# Encoding of natural variability in a dexterous motor skill over multiple days in the cortex of freely behaving mice

**DOI:** 10.1101/2024.12.20.629771

**Authors:** Elizabeth A. de Laittre, Jason N. MacLean

## Abstract

Skilled, goal-directed movements exhibit trial-to-trial variability even in experts, particularly in response to dynamic environmental conditions or when perfect repetition is not required for success. Identifying where, to what extent, and how stably this variability is encoded in the nervous system is essential for understanding how learned movements are robustly maintained over time yet flexibly executed on each trial. We record calcium fluorescence activity in forelimb motor cortex (M1), a key node in the multi-areal network responsible for movement control, in freely-moving mice of both sexes as they performed a self-paced, precision reach-to-grasp task. High trial counts and rich single-trial variability enable rigorous statistical analysis of moment-to-moment movement encoding across matched behavioral sets over five days. Approximately 80% of recorded neurons significantly encoded paw, digit, and head movements during reaching, as quantified using linear models. Across days, encoding similarity shows a small but measurable decline that increases with the interval between recording sessions. This drift is heterogeneously distributed across the population, with many neurons retaining high encoding similarity even in sessions four days apart, as assessed using shuffle controls and comparison to encoding for trial-averaged movements. Thus, over the timescale examined, M1 is capable of maintaining stable encoding of movement details at the level of single cells, even for complex, sensory-guided tasks like reach-to-grasp. Together, these results raise the question of whether downstream circuits support consistent behavior by preferentially relying on neurons with greater stability or instead through population-level readout that is robust to a modest level of representational change.

**Significance Statement:** Skilled actions must be both precise and flexible, adapting to changing situations and environments. How the brain achieves consistent performance when the circumstances of each attempt rarely repeat remains poorly understood. By tracking hundreds of neurons in mouse motor cortex during a precision reach-to-grasp task, we show that individual cells stably encode fine details of paw, digit, and head movements across multiple days. This stability provides a reliable foundation for consistent execution while supporting adaptability in real-world contexts. Our findings indicate that motor cortex integrates postural information with skill-specific forelimb and digit signals, revealing how single neurons support robust yet flexible motor control, a capacity that remains a major challenge for artificial systems.

## Introduction

Even well-practiced behaviors exhibit variability with each attempt. In basketball, for instance, no two shots a player makes are identical. Each is shaped by factors such as starting posture, fatigue, and interactions with defenders and teammates. While generating goal-directed movements under such conditions remains a significant challenge in robotics (Billard and Kragic, 2019), biological nervous systems excel at this. To understand how motor skills are consistently executed across varying contexts, it is crucial to determine how movement variability is represented in neural circuits and whether the same activity patterns generate the same skill across days. This will clarify the circuit-level precision required to adapt to perturbations while maintaining reliable performance.

Prehension, the act of reaching and grasping objects, is one such learned behavior that is tractable for study in animal models and conserved across taxa (Sacrey et al., 2009; Whishaw et al., 1992; Guo et al., 2015; Becker, Calame, et al., 2020; Whishaw et al., 2017; Aparicio et al., 2026). In mice, motor cortex, specifically caudal forelimb area (CFA), plays a central role in the brain-wide circuit controlling reach-to-grasp and other precision reaching behaviors (Guo et al., 2015; Wang et al., 2017; Miri et al., 2017; Galiñanes et al., 2018), which rely on online, active control rather than pre-planned, ballistic movements (Becker, Calame, et al., 2020). CFA has recently been shown to encode low-level details of forelimb movement such as muscle activation and kinematics, e.g. joint position and velocity (Omlor et al., 2019; Koh et al., 2025; Grier et al., 2026). However, in all such studies, mice were head-fixed, reducing the need to coordinate the forelimb with body posture and altering reaching submovements (Whishaw et al., 2017).

In freely-moving mice, body position and posture constrain the forelimb and digit movements required for a successful grasp. Thus, freely-moving reach-to-grasp requires coordination of the whole body, from gross limb and trunk movements to fine digit control, guided by rapid sensory feedback processing. Although coordinated digit use and posture-related activity in rodent sensorimotor cortex have been reported (Whishaw et al., 2017; Barrett et al., 2020; An et al., 2026; Mimica et al., 2018; Disse et al., 2023), most studies of reaching in rodents have focused on proximal forelimb (elbow and shoulder) kinematics. Clarifying whether and how CFA integrates information about multiple coordinated body parts to guide the forelimb during reaching is essential to understand its role in this complex, naturalistic behavior.

Investigating how neural encoding of movements changes over time can reveal properties of plasticity mechanisms subserving motor skill learning and maintenance (Micou and O’Leary, 2023; Driscoll et al., 2022). However, the stability of moment-to-moment encoding for low-level movement features, such as joint position and velocity, remains underexplored. To overcome neuron turnover across recording sessions, one study in rhesus macaques aligned low-dimensional population activity, finding that movement details can be decoded stably across sessions (Gallego et al., 2020). Whether these population “modes” reflect stable single neuron activity remains unclear. In cases where cells are tracked over days with high confidence via calcium imaging or continuous electrophysiological recording, studies have found day-to-day consistency of trial-averaged neural activity associated with highly stereotyped movements (Jensen et al., 2022; Katlowitz et al., 2018; Aparicio et al., 2026; but see Liberti et al., 2016). It is unclear whether such consistency extends to moment-to-moment encoding during flexible movements that require sensory-driven adjustment.

To address these gaps, we combine high-speed-video-based movement reconstruction with head-mounted one-photon calcium imaging in mice performing a whole-body, dexterous reach-to-grasp task that yields high trial counts with significant trial-to-trial variability. Despite this within-session variability, individual mice consistently explore a similar movement space across days, enabling matched datasets to directly compare encoding of multiple movement variables—kinematics of the head, forelimb, and digits—across days. We find that single-trial variance of these movements is encoded in mouse forelimb motor cortex and this encoding remains highly stable across days at the single-cell level, supporting reliable yet flexible skill execution over time.

## Methods

### Mice

All experimental procedures were approved by the University of Chicago Institutional Animal Care and Use Committee and were performed in compliance with the Guide for the Care and Use of Laboratory Animals. Data were collected from B6.Cg-Igs7tm148.1 (tetO-GCaMP6f, CAG-tTA2) Hze/J mice (Jax strain 030328, also known as Ai148D) of either sex (n = 2 female, n = 3 male). The line was originally obtained from Jackson Labs and bred in house. Mice were between 10 and 12 weeks old at the start of the shaping and training protocol.

### Surgical procedures

Animals underwent three surgical procedures: AAV-Cre injections, prism implantation, and baseplate implantation. For all surgeries, anesthesia was induced with isoflurane (induction at 4%, maintenance at 1–1.5%). During AAV injection, two small burr-hole craniotomies were drilled to allow insertion of a micropipette and injection of virus. Injections were made at three depths (650, 400, and 200um below the surface) at two locations targeting caudal forelimb area (0.25mm anterior and 1.25mm lateral to bregma, and again at 0.25mm anterior and 1.75mm lateral to bregma). After injections, the burr holes were sealed with dental cement and the skin sutured. Animals recovered for a minimum of seven days before prism implantation. During prism implantation, a 2mm diameter craniotomy was drilled, centered at 0.25mm anterior and 1.5mm lateral to bregma. To ease insertion of the prism into the cortical tissue, a stereotaxically mounted scalpel was used to make a 1.25mm wide incision running along the medial lateral axis (parallel to the sagittal plane), centered at the same coordinates as the craniotomy; the incision ran to a depth of 800um. The prism was then inserted into the tissue along this incision to the same depth, with the angled part of the prism directed posteriorly. The resulting field of view is thus anterior to the site of insertion. The remaining surface of brain tissue was covered with Kwil-sil (World Precision Instruments) and the prism was cemented in place. Animals recovered for a minimum of seven days before baseplate implantation. During baseplate implantation, a plastic baseplate was cemented above the prism and grin lens, placed to allow the best alignment of the microscope with the focal plane of the prism. This final procedure is non-invasive, requiring only the application of cement to previously implanted hardware; there is no contact with soft tissue. Animals recovered for a minimum of one full day before the start of food restriction and habituation in the arena.

### Behavioral training

The entire behavioral experience consisted of three parts: habituation, shaping, and training. During all phases, the mice were briefly (∼1min) and lightly anesthetized with isoflurane so that the miniscope could be attached to their headplates. Mice were then placed in the arena and allowed to recover completely (defined by showing unencumbered, normal ambulatory movement, at least 5 minutes) before the behavioral protocol for that day began. The arena was 15cm x 15cm with a reach port (6mm wide rectangular opening that ran from the floor up ∼8cm) located in the center of one of the walls (see schematic in Fig. 1B). The wall with the reach port was slanted away from the arena at a 30 degree angle relative to vertical to enable the mice to maneuver better at the reach port while wearing the miniscopes on their heads. Removable walls were placed on either side of the reach port to limit possible body positions around the reach port to make the pedestals reachable with only the left paw during the early phases of shaping and training (see schematic in Fig. 1B). For some animals these walls were present throughout training, including the days used in this study; for other animals, these walls were removed before the days included in this study. There was no notable impact on the overall set of body positions these animals used in a given session before or after removal of the walls.

**Figure 1:**
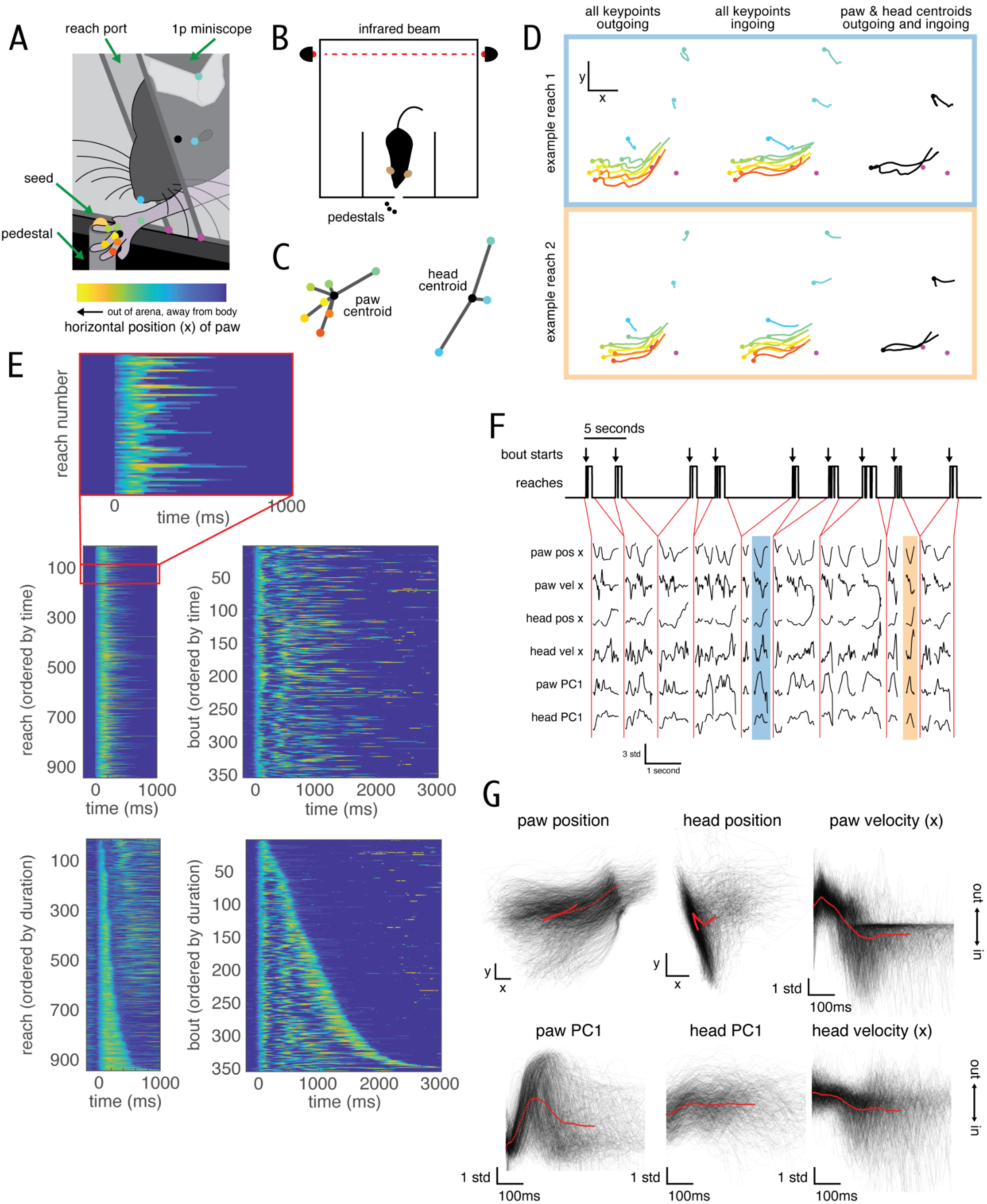
A dexterous reach-to-grasp task elicits variable yet spatiotemporally coordinated forelimb, digit, and head movements. A) Closeup view of freely-moving mouse reaching for a millet seed reward, with 10 tracked keypoints on the paw and head marked, from the viewing angle of the tracking camera. Colorbar at bottom indicates paw position in x relative to the view shown here; relevant for the heatmaps in E. B) Schematic of freely-moving mouse in arena viewed from above, indicating location of pedestals, reach port, and infrared beam. Arena floor dimensions are 15cm by 15cm. C) Schematic of tracked keypoints on the paw and head for the keypoint locations shown in A, and distances to the paw and head centroids (black dots) shown in gray. Colors match those in A. Paw and head dot locations are not drawn to the same scale. D) Two example reconstructed reaches. Colors match those in A and C. For all panels, dots indicate the endpoint (point of the reach that is furthest from the arena). Left column, all paw and head keypoints during the outgoing part of the two reaches. Middle column, same for the ingoing part. Right column, the paw and head centroids only for both the outgoing and ingoing parts of the reaches. Magenta dots indicate the base corners of the reach port, as in A. E) Paw position in the x dimension plotted for all reaches and bouts in one daily session for an example mouse (mouse 5). Colormap is under the schematic in A; yellow colors indicate the paw is further outside the arena, blue colors indicate closer to or inside the arena. Top, reaches sorted in the order in which they occurred in the session (left), and the same reaches resorted into bouts, again in the order in which they occurred (right). Bottom, the same reach and bout data sorted according to reach and bout duration, respectively. F) A subset of kinematic variables for some example reaches, including the two example reaches in D indicated by blue and orange shading. Top, binary variable indicating when the animal is reaching (paw is outside the arena; high indicates reaching, low indicates not reaching), with an arrow at the start of each bout. Bottom, time courses of 6 of the 14 kinematic variables extracted via keypoint tracking, during the bouts plotted above. G) Reaches show measurable trial-to-trial variability in all extracted movement features, enabling the study of neural coding for these continuous variables. All reaches in one daily session for an example mouse (mouse 5) for the same 6 variables as in F. Individual reaches are plotted in transparent gray, the average (time-locked to reach onset) is plotted on top in red. Scale bars for paw and head position correspond to 30 pixels in x and y; the view is the same as is schematized in A and for the example reaches in D. Scale bar for paw velocity, head velocity, paw PC1, and head PC1 are 100ms and 1 standard deviation of each variable.

#### Food restriction, Habituation and Shaping

At least one full day after baseplate implantation, mice began food restriction, in which they received a limited amount of pellet food, titrated to maintain their weight between 85% and 90% of their baseline weight. Mice were food restricted for at least one week before habituation began, sometimes longer in order to achieve a weight between 85% and 90% of baseline. Mice were singly housed throughout food restriction, habituation, and training.

Mice were habituated to the training arena and the miniscope for at least seven days before shaping began. During habituation, mice were placed in the behavioral arena for twenty to thirty minutes, once per day. At the start of habituation, twenty millet seeds were placed on the ground inside the arena next to the reach port, to encourage an association between the reach port and food.

During shaping, mice were again placed in the behavioral arena for twenty to thirty minutes, once per day. Instead of being presented inside the arena, seeds were presented in two places outside the arena: approximately ten seeds were presented on the ground outside the arena immediately in front of the reach port. These were reachable by the animal’s tongue. Seeds were also presented in a low square trough outside the arena near the reach port. The trough was positioned a few millimeters laterally to the center of the reach port, such that the animal would only be able to successfully reach the trough with its contralateral forelimb. All mice were shaped and trained to use their left paws. The goal of shaping was to train the animal to associate reaching its forelimb outside the arena with getting food. Functionally, during shaping, the mice would scoop or knock seeds out of the trough and onto the ground, then lick them up with their tongues. Any large magnitude forelimb movement through the reach port was considered a reach. In order to move from shaping to training, mice needed to make at least 50 reaches with their left forelimb during a twenty minute period within the shaping session, and at least 80% of their total reaches in that period needed to be with their left forelimb. This shaping procedure is based on that described in Chen et al. (2014). Shaping was performed daily until mice achieved these criteria within a single shaping session. Often during the last shaping session before training, mice far exceeded these criteria, reaching 75-100 times with their left forelimbs. Mice began the training protocol the next day after achieving the shaping criteria. If, on the first training day, the animal did not show interest in reaching for the seeds on the pedestal, or reached very few times, the training session was converted back to a shaping session, to reinforce the association between using their forelimbs and getting seeds. The animal was held to the same requirements as before for returning to training.

We found that on the first day of shaping there is a critical moment requiring manual involvement. Sometimes at the beginning of shaping, the first reaches that mice make out of the port do not make contact with the seed trough or do not make strong enough contact to knock any seeds out. We observed that many mice give up reaching entirely after only a few tries if their initial reaches do not lead to reward. We found that if we reward the first one or two left-paw reaches by manually knocking seeds out of the trough immediately after the reach, the animals continue to reach and quickly learn the desired association between left-paw reaches and food reward.

#### Training

During training, seeds were only presented to the mouse on the seed delivery pedestals. Pedestals were positioned relative to the reach port such that they were within reaching distance with the paw, but out of reach of the tongue; this positioning was adjusted for each mouse because of size differences across mice, particularly across gender. The pedestals were narrow such that they could only hold one seed each and the seed was easy to knock off and out of reach. On a given training day, the mice were in the arena for between 75 and 90 minutes. Periods of time where seeds were presented (and mice subsequently reached) were interleaved with break periods during which no seeds were presented. Seed presentation periods lasted 3-5 minutes, followed by 2-4 minutes of break. There were 8-12 seed presentation periods per session, totaling 40 minutes of seed presentation (and subsequent reaching behavior) per day. Training was performed at approximately the same time each day for a given mouse. Imaging took place every training day. For each mouse, five sessions from late in training were used in this study, starting between day 20 and 22.

#### Seed delivery

There were two different kinds of seed delivery methods used for the mice included in this study. Mice only ever experienced one seed delivery method throughout their training.

For two of the mice (mice 1 and 2), there were three permanent seed delivery pedestals positioned in an arc outside the reach port approximately equidistant from the close edge of the reach port and referred to as medial, lateral, and far lateral (schematized in Fig 1B). Their positioning was offset to the mouse’s right such that the pedestals were not reachable with the right forelimb. Seeds were delivered manually by the experimenter. Mice were required to turn away from the seed port and walk towards the back of the arena in order for a new seed to be delivered, thus initiating a new “trial”. Only one seed was placed per trial. The order of seed placement was random but counterbalanced such that there was approximately the same number of deliveries per pedestal in a given session.

For one of the mice (mouse 3), the same manual delivery method was used with the three permanent pedestals, but seeds were only placed on the center pedestal, referred to as the lateral pedestal.

For the final two mice (mice 4 and 5), there was a single mobile seed delivery pedestal which was moved laterally between two different positions throughout the session by a linear actuator. Their positioning was offset to the mouse’s right such that the pedestals were not reachable with the right forelimb. The two pedestal positions are a subset of the three positions used for manual delivery (schematized in Fig 1B), using only the two pedestals closest to the mouse’s midline, referred to as the medial and lateral pedestals. For these mice, seeds were delivered automatically: the pedestal was lowered by a linear actuator into a large trough of seeds and then lifted again, having caught a seed. The design of our automated seed delivery system is very similar to that described in Ellens et al. (2016) using the pellet hopper and delivery arm, but without a reaching shelf. The trough of seeds was multiple centimeters below the floor level of the arena and thus out of reach. The schematic in Fig 1A is based on our automated seed delivery system. The order of the two delivery positions was random in a given session, determined by a Bernoulli random variable in custom arduino code. To trigger the delivery of another seed, mice had to break an infrared beam at the back of the arena.

For the manual seed delivery, if mice turned away to trigger another seed delivery but the previous seed was still present on the pedestal, the seed was left in position for up to two additional trials before that seed was removed and a new seed was placed.

For the automated seed delivery, there was no timeout for unsuccessful reaches and no requirement that the seed be removed from the pedestal before triggering another seed delivery.

For mice that had seeds delivered to multiple pedestal locations, they were trained on all pedestal locations from the beginning of training.

### Behavioral video recording and kinematic reconstruction

#### Videography and keypoint extraction

We took high speed, color video of the mouse during reaching at a frame rate of 200Hz using a Flir 16S2C USB3 Blackfly color camera (BFS-U3-16S2C-CS). All kinematic data used in this paper is captured in 2D, with a camera that is at an approximately 30 degree angle from head on, offset to the mouse’s left to best view the left forelimb and paw. Additionally, the camera view is angled approximately 20 degrees from above. The schematic in Fig 1A represents the camera viewing angle. We used DeepLabCut to extract a variety of kinematic features from this video data (Mathis et al., 2018; Nath et al., 2019). On the paw, we tracked the joints of the first three digits and the wrist; on the head, we tracked the nose tip, corner of the eye, and a prominent corner of the surgical implant (Fig. 1A). After extracting keypoints using DeepLabCut, kinematic data was further processed with custom MATLAB software. Points that were identified as poorly labeled (because of large distances traveled from the previous timepoint, or because markers that are anatomically close together were labeled as too far apart) were linearly interpolated over. Data was then smoothed: missing frames were linearly interpolated over, we smoothed using a butterworth filter (4th order, low-pass filter with 50Hz cutoff), then removed the interpolated (originally missing) frames.

#### Alignment across days

The position of multiple non-moving features of the arena (referred to as static markers) were extracted from the videos and used to align kinematic data across days. We adjusted for translation only: the x and y position of the most reliably tracked static marker was subtracted from all the kinematic variables. Generally this translation was minimal because the cameras were fixed in place relative to the arena. We also extracted the position of the pedestal and the presence of a seed for each reach automatically from the video data.

#### Calculating kinematic variables of interest

We calculated the centroid of the paw as the average position of seven markers on the paw (wrist, digits 1-3 bases and middles). The centroid of the head was calculated as the average of all three markers on the head (nose, front corner of the eye, and corner of the microscope). See Fig 1A and C for visualization of these markers. To calculate a low dimensional estimate of paw shape and orientation, we subtracted the paw centroid position from the seven paw markers, in x and y separately, then performed pca on these 14 dimensions (seven markers, in x and y). We projected data onto the first 4 dimensions, then included it in the linear model. We performed the same procedure for the three markers on the head to capture an estimate of head orientation. We performed pca on the 6 original dimensions and projected data onto the first 2 dimensions to include in the linear model. We pooled aligned data across all five days before performing PCA, such that the PCA values were meaningful relative to each other across days. For one mouse (mouse 3), PCs of the paw markers were not included in analysis due to poor digit tracking.

Note that our kinematic data is two dimensional and extracted from a single camera view, such that the head and paw PCs capture mixtures of rotation and translation as well as shape. The position of the paw and head in 2D are subtracted out from paw and head markers, respectively, before performing PCA to remove the effects of translation in x and y on our estimates of paw and head rotation and shape. However, the translation in z (depth, towards and away from the camera) cannot be subtracted out and thus is captured in our PCA. To minimize the impact of translation in z on our ability to capture paw orientation and shape from a 2D image, we aligned the camera such that the major axis of paw movement is parallel to the imaging plane. Hence, we refer to the paw PCs as capturing orientation and shape of the paw.

We calculate velocity from the position data using the following approach: we first calculate the difference between position in consecutive timepoints and divide that by the amount of time between timepoints (generally 5ms). We then treat that as the velocity at the halfway point between the observed timepoints and use linear interpolation to infer the velocity at the observed timepoints.

#### Interpolation

For the linear encoding model analysis, kinematic data (200Hz) was interpolated to the timepoints of the neural data (30Hz) using the MATLAB function interp1.

#### Segmenting reaches

Reaches are defined using the paw centroid position relative to a static marker on the reach port (right magenta dot in Fig 1A): the paw was considered outside the arena when it was to the left of the static marker, and inside the arena when it was to the right of it. The onset of a reach was identified as the first video frame where the paw centroid was outside the arena and similarly the end of the reach was identified as the last video frame when the paw centroid was outside the arena before it re-entered. By this definition, reaches often consist of one out and back motion, but could include multiple out and back motions if the paw centroid did not pass to the right of the static marker in between.

Reaches were clustered into multiple reaches in quick succession, followed by a longer break during which the animal left the reach port to trigger the delivery of another seed (see Fig 1F top for illustration). We refer to such a cluster as a bout. Bout starts were identified using the inter-reach interval: an inter-reach interval of greater than 1 second separated two bouts. For some sessions for some mice, we used a shorter threshold between bouts (as low as 700ms), depending on the distribution of inter-reach intervals and visual inspection of the behavior.

All reaches to all pedestals were included in the neural coding analyses, since all reaches constituted movements that can be used to study the motor cortical encoding of different kinematic features.

#### Performance on the task and calculation of success rate

A successful reach is one where the animal grasped the seed and brought it into the arena. Success rate is defined as the proportion of reaches that are successful out of all reaches made with a seed present on the pedestal in the medial or lateral position. Reaches where a seed is not present at the beginning of the reach are not included in the total. Success was scored manually by watching the video recording of each reach.

For all animals, only reaches when the seed was on the medial and lateral pedestals were included to calculate the success rates. Of the two animals trained with seeds presented on the far lateral pedestal, neither ever successfully retrieved a seed from that pedestal, and it was deemed too far for them to reach from the port. For these animals, reaches while a seed was presented at that position were not included in success rate calculations but were included in neural encoding analyses.

#### Inclusion of reaching data in analysis

Successful versus failed reaches: Due to the nature of the task, successful reaches naturally exhibit lower variance across multiple kinematic features compared to failed reaches. For example, the paw must pass within a certain distance of the pedestal during the reach for the animal to grasp the seed. In all neural coding analysis, both PETHs and linear models, we include both successful and failed reaches, as both sets of movements can be used to study the motor cortical encoding of different kinematic features. This benefits our analyses because it both increases the sample size of reaches and timepoints, and it also provides a wider range over which to quantify the neural encoding of each movement variable.

Seed position: For the same reasons described for successful and failed reaches, we also included in our analyses all reaches regardless of the position of the seed (see Methods section Seed Delivery for description of seed presentation positions and Fig. 1B for schematic)

Timepoints during reaching: We had the least occlusion and the most reliable tracking of the paw and digit markers during reaching (when the paw was outside of the arena) so we focus our analysis of neural coding on the animal’s movements there.

Hence, only timepoints during reaching were included in the linear model analyses. Timepoints while the paw was inside the arena, including when the animal was eating, grooming, and locomoting to trigger seed delivery, were excluded from linear model analyses.

### One-photon calcium imaging

#### Image acquisition

One-photon calcium imaging was performed using an Inscopix nVista 3.0 miniscope (excitation centered at 475 +/- 7 nm, collection centered at 535 +/- 25 nm) controlled by Inscopix Data Acquisition Software (IDAS). A baseplate was chronically attached to the animal’s skull and centered over the chronically implanted grin lens and prism (see *Surgical procedures*). The lightweight (2.2g) miniscope was attached to the baseplate during recording sessions. Data was collected from a single imaging plane at 30Hz, referred to as the field of view (FOV) for this mouse. Each recording session lasted 40 minutes, broken up into 3 to 5 minute recording chunks, each separated by a break of anywhere from 2 to 4 minutes. Imaging was continuous during a recording chunk, but paused during breaks. When analyzing imaging data from a given session, data from all chunks in that session were concatenated and analyzed together.

The field of view corresponds to approximately 900 x 650 micrometers at the plane of focus, captured in 1280 x 800 pixels. Use of the 45-degree prism means that our field of view is perpendicular to the surface of cortical tissue, providing a coronal rather than axial (horizontal) imaging plane. The cortical surface was apparent at the top of each individual FOV. Based on the size of the FOV and distance from the surface, our cell sets include predominantly, if not exclusively, layer 2/3. For mice with a C57BL6 background, the thickness of motor cortex for coordinates within the caudal forelimb area has been observed to be 1.378 ± 0.021 mm, (*n* = 11 mice; Wood and Shepherd, 2010). Previous characterizations of the laminar structure of mouse motor cortex find that the layer 5a boundary with layer 2/3 occurs at a normalized depth of 0.40 ± 0.02 from the pia (*n* = 3 mice), with a normalized depth of 0 corresponding to pial surface and 1 corresponding to white matter (Oswald et al., 2013). The layer 2/3–layer 5a boundary thus occurs approximately 550 micrometers below the pial surface. Given that the vertical extent of our field of view is up to 650 micrometers and that the prism was implanted such that the pial surface is visible at the top of all FOVs, our data includes predominantly cells in layer 2/3, potentially with some cells from superficial layer 5a at the very ventral extent.

#### Locating the same field of view over days

The same field of view was imaged across all days of training. Because the prism is chronically implanted into the neural tissue, the only degree of freedom in identifying the field of view is depth (because of the 90 degree rotation from the prism and its orientation, more “shallow” corresponds to more posterior, whereas deeper into the tissue corresponds to more anterior). On the first day of training, the depth with the most active cells was selected while the animal was moving around the arena but not performing the task. Then, some small blood vessels were identified and used as fiduciary markers to find the same plane on subsequent days.

Imaged FOVs are estimated between 50 and 100 micrometers deep in the tissue from the surface of the prism.

#### Pre-processing of calcium imaging data and region of interest (ROI) extraction

Data was pre-processed using the Inscopix Data Processing Software (IDPS) MATLAB API and included 4x downsampling by averaging (a 4x4 square of pixels is averaged to one pixel), spatial filtering by frame (bandpass with low cutoff 0.005Hz and high cutoff 0.5Hz, where Hz refers to units of 1/pixel), and across-frame motion correction using TurboReg (Thevenaz et al., 1998).

We exported the resulting stacks of images for cell identification, and used Caiman CNMFE (Zhou et al., 2018; Giovannucci et al., 2019) to extract spatial footprints and single-cell fluorescence time series. CNMFE is an extension of CNMF (Pnevmatikakis et al., 2016) designed specifically for one-photon endoscopic calcium imaging data. The algorithm includes a more complex background model designed to capture fluctuations due to out-of-plane cells and neuropil, with the goal of removing these fluctuations from the algorithm’s ROI timeseries output. For more details, please see Zhou et al., 2018 and Giovannucci et al., 2019. For each cell separately, we convolved its inferred spike data (S output from Caiman) with a gaussian of 30ms standard deviation. This is the neural activity signal used for all analysis unless otherwise noted.

Due to computational limitations, each 40 minute session was processed via Caiman in batches. Data from a session was broken up into 3 batches of approximately equal size by recording chunk (see previous methods section *Image acquisition*), distributing the recording chunks such that each batch contained neural data from the beginning, middle, and end of a session (e.g. batch 1 contained recording chunks 1, 4, 7, etc.). After passing each batch of imaging data through the Caiman CNMFE algorithm, we matched cells across batches using CellReg (Sheintuch et al., 2017), with the requirement that weighted spatial footprints had to have a spatial correlation of 0.9 across batches to be considered the same cell. Only cells found in all batches were included in the final cell set for a given session. The spatial footprint for a given cell for a given session was computed by averaging across its spatial footprints from each batch. Time series variables (Caiman variables C, C_raw, and S) for each cell were z-scored within batch before de-interleaving and concatenating across batches.

#### Registering cells across days

We registered cells across days using CellReg (Sheintuch et al., 2017), with a strict, fixed requirement that across two consecutive days, weighted spatial footprints had to have a spatial correlation of 0.9 to be considered the same cell.

#### Reach modulation via ANOVA

Reach modulation was determined for each cell individually via an ANOVA comparing its activity during reaching timepoints with its activity during all other timepoints, which include other behaviors such as locomotion, grooming, food manipulation, and food consumption. Significance of reach modulation was assessed with a p-value of 0.05 with Bonferroni correction for the number of cells in the session (that day’s recording session for that mouse).

### Peri-event time histograms (PETHs)

We calculated PETHs by averaging neural activity in each time bin across trials. Each reach is one trial. Neural data from all reaches were included in PETH calculation, regardless of success or seed position. Here, t = 0 indicates the start of the reach, also referred to as reach onset. For visualization, PETHs were normalized such that the lowest value = 0 and highest value = 1 within the shown time window, [-500 1000]ms relative to reach onset. For comparing PETHs over days, PETHs were normalized in the same way but within a narrower window. This shorter window was used when calculating the correlation between PETHs for the same cell found on different days. For PETHs locked to reach onset, the window was [-100 500]ms relative to reach onset (where reach onset is at t = 0); for PETHs locked to bout onset, the window was [-300 1500]ms relative to bout onset (where bout onset is at t = 0).

#### Cell ID shuffle control

The cell ID shuffle distribution was generated by calculating the correlation between PETHs for non-same cells found on different days. For each pair of days (n = 5 days, yielding n = 10 day-pairs), we randomly sampled a cell from each day, ensured they were not registered as the same cell, and computed the correlation between their PETHs. We repeated this process 1000 times for each day-pair, yielding a total of 10,000 samples in the shuffle distribution. Shuffle distributions were generated separately for each mouse. This control distribution is similar to the one used by Jensen et al., 2022. It yields the correlation values expected if on any given day, each neuron’s trial-averaged activity pattern is randomly sampled from the distribution of available patterns in the population. Thus this distribution represents complete instability of encoding at the individual neuron level, but with constant activity statistics at the population level.

### Linear encoding models of single neurons

On each day separately, we fit a linear model where we predicted the activity of each cell based on measured behavioral and task variables. We used an instantaneous encoding model: neural activity at time t was predicted from behavior and task variables at time t. No time-shifted versions of any predictors were included. Behavioral variables were all interpolated from the original 200Hz to the 30Hz framerate of the neural data (see *Interpolation* section in *Behavioral video recording and kinematic reconstruction*). Here and throughout this work, the data for a given session refers to the neural and behavioral data from one mouse from one day. There are 25 sessions total: 5 mice, with 5 days for each mouse.

#### Data inclusion and predictor sets/design matrices

For each mouse, data from five sessions from late in training were used in this study, starting between day 20 and 22 (see *Training*). In total, 15 predictor variables were included: the position and velocity in x and y of the centroids of the paw and the head (8 variables), the first four principal components (PCs) of the paw markers, the first two PCs of the head markers, and time within the session (referred to as the time-in-session predictor). Time-in-session does not include breaks between recording chunks, meaning the variable is equivalent to the index of the calcium imaging data; it increases monotonically over the course of the session.

Time-in-session is included as a predictor to capture slow, session-wide changes in neural activity due to changes in behavioral state, internal state, and task engagement reflecting variables such as fatigue, satiety, arousal, motivation. All linear model results were repeated excluding the time-in-session predictor; results are included in Supplementary Figure 5C-J and Table S1. For one mouse (mouse 3), PCs of the paw markers were not included due to poor digit tracking. Only timepoints during reaching (when the paw was outside the arena) were used to fit the linear models. Timepoints while the paw was inside the arena, including when the animal was eating, grooming, and locomoting to trigger seed delivery, were excluded from linear model analyses (see next section and Supplementary Figure 4D for the number of timepoints per session broken down by mouse and day). As with the PETHs, all reaches were included in this analysis, regardless of success or seed position.

#### Covariate balancing across days to allow for direct comparison of linear fits

In order to directly compare properties of our linear model fits across days, we must ensure that each predictor has a similar distribution to itself over days. We must also ensure that the joint distribution of predictors is similar over days, which captures the correlations between predictors. Without this step, we cannot guarantee that any observed encoding instability is not a result of the animals exploring a slightly different part of kinematic space each day. We subsample from each day’s collection of reach timepoints in the following way to create a similar distribution for each day within the high-dimensional space of all predictors.

Because we want to compare the encoding for the same absolute values of the predictors, we first z-score on all days together rather than on each day separately. For each mouse separately, we treated each predictor as follows: We concatenated all reach timepoints for that predictor across all five days, then z-scored. We then separated this data back into days and performed the covariate balancing procedure described below.

We use a greedy matching algorithm without replacement to balance the joint distributions of all behavioral predictors across days. In a given day’s joint distribution, each sample is a reach timepoint; it corresponds to a vector of all kinematic predictor values at that timepoint (14D for mice 1, 2, 4, 5, and 10D for mouse 3). The time-in-session predictor is not included in this vector during covariate balancing. We use the day with the fewest reach timepoints as the reference day. For each timepoint in the reference day, we identify the timepoint in each other day that is most similar to the reference day timepoint. Similarity is calculated as the correlation between the vectors of kinematic predictors. Timepoints on the reference day are matched sequentially. After a timepoint on a non-reference day is matched to a reference day timepoint, it is no longer available for matching to later reference day timepoints; hence, the algorithm is greedy. After finding matches for all reference day timepoints, we remove matches in two phases. First, we remove non-reference day timepoints that have a correlation with their matched reference day timepoint that is below 0.9. There may be a different reference day timepoint for which this timepoint has a correlation greater than 0.9, but the algorithm guarantees that reference day timepoint already has a better match on the given non-reference day. Second, we remove reference day timepoints (and all their associated matched non-reference day timepoints) that have fewer than 2 remaining matches, meaning that reference day sample has only one other day where a sufficiently similar timepoint was found. Because the algorithm is greedy (steps through the timepoints sequentially, which prioritizes good matches for timepoints seen earlier in the algorithm), we randomize the order that the reference timepoints are matched in; this means that the best matches are not biased towards the beginning of the session, but instead distributed throughout.

If an animal’s behavior is dissimilar over days, with less overlapping predictor sets, the covariate balancing procedure would yield a very small number of samples left to fit the models with. Even before subsampling, the probability density functions of our predictor sets are largely overlapping, indicating very similar behavior across days. Thus, covariate balancing leaves us with a large number of samples for each session to fit the models with (12889 +/- 4933 [7276 22175] samples per session (mean +/- st. dev. [range]), which constitutes 68.7% +/- 17.2% [31.5% 95.%] of timepoints in the original sessions (mean +/- st. dev. [range]); Supplementary Figure 4D). Sessions where lower percentages of timepoints were kept generally constitute days where the animal performed a higher number of reaches but with consistent distributions of behavioral features, such that subsetting is matching the number of timepoints across days rather than to removing dissimilar behavior (for probability density functions for an example predictor for all days for an example animal, see Supplementary Figure 2A).

After covariate balancing, each predictor is again z-scored within day before fitting the models. All single day linear model results presented in this paper are fit with predictor sets that have been covariate balanced across all five days for that animal.

#### Assessing the impact of covariate balancing on predictor distribution similarities

To assess whether the similarity of predictor distributions across days changed as a result of the covariate balancing procedure, we compared the distributions using the non-parametric similarity metric Wasserstein distance (Earthmover’s distance). Wasserstein distance quantifies how much distribution mass needs to be moved multiplied by how far in order to make two distributions equivalent.

We compute Wasserstein distances for each predictor separately. Before computing these distances, we z-score each predictor. Wasserstein distance is sensitive to the absolute value of the predictors, which varies greatly from predictor to predictor because they are capturing very different physical properties (e.g. position versus velocity). Z-scoring allows us to meaningfully compare distances for different predictors. Z-scoring for a given predictor is performed on all 10 distributions together (the before-balancing distribution and the after-balancing distribution for all five days). Thus the absolute values of the predictors are kept meaningful relative to each other, and Wasserstein distances between before-balancing distributions can be directly compared to distances between after-balancing distributions. Finally, we normalize each distribution by total probability to yield a probability density function (PDF). Normalizing to a PDF ensures that Wasserstein distances do not simply reflect the fact that the number of samples in the distributions are more similar across days after balancing than before.

Wasserstein distances are then computed between before-balancing PDFs for all pairs of days (5 days choose 2 = 10 day-pairs), and similarly for after-balancing PDFs. There are 10 before balancing distances and 10 after balancing distances for each predictor for each mouse. To compute the grand mean Wasserstein distances for each predictor before and after balancing (Supplementary Figure 2C), we first average across the 10 day-pairs to yield a single mean distance value for each predictor for each mouse, then we average across the 5 mice.

#### Fitting the single-trial family of models

Single-trial models refers to the standard instantaneous linear model fit where we predict neural activity at time t from behavioral and task variables observed at time t. Models were fit using ridge regression, using the MATLAB function ridge() with a ridge parameter equal to the number of timepoints (samples) in that day; see previous methods section *Covariate balancing across days to allow for direct comparison of linear fits* for statistics on the number of timepoints per day.

We fit a family of models for each cell on each day. Full models were fit using all 15 predictors specified in the methods section *Data inclusion and predictor sets/design matrices*. As described in the main text, we also fit a single variable model for each predictor that included only that predictor, as well as a leave-one-out model for each predictor that included all predictors except for the one in question. Delta-R-squared values are computed as the drop in R-squared from the full model to the leave one out model for the predictor in question, as schematized in Fig 5a.

We used a train-test split of 70% train, 30% test. We did the following procedure to ensure that each train-test split contained a balanced sampling of timepoints from across the session. For all the timepoints included in a day for a given mouse, we split the data into 100 chunks. We then grouped every 10th chunk together, resulting in 10 sets. For example, chunks 1, 11, 21, etc. went into set 1, while chunks 2, 12, 22, etc went into set 2. We then chose 7 random sets to use as the train set, and held out 3 chunks as the test set. We repeated this procedure for 10 cross-validations for all model fits. Within a day, the train-test splits for each cross-validation were the same for all cells.

All reported R-squared and delta-R-squared values are averaged across 10 cross-validations of the train-test split.

Delta-R-squareds are calculated within each cross-validation, then averaged across cross-validations. Note that due to the train-test split, R-squared values can sometimes be negative. Similarly, the delta-R-squared (leave one out models) can sometimes be positive, indicating that removing the predictor in question improved the test R-squared values. This typically occurs for small R-squared values or poor fits, where the model is fitting to noise in the train set and thus cannot generalize to the test set.

#### Significance testing of full model R-squared values

To test whether a full model fit was better than expected by chance, we compare each cell’s full model R-squared value to a timepoint shuffle null distribution. For each cell, we generate a null distribution by shuffling the timepoints of the neural activity (predicted variable) and refitting the full model for that day. We repeated this procedure 2,000 times per cross-validation, for a total of 20,000 shuffles. If the observed, cross-validated full model R-squared value was greater than a threshold percentile of time-shuffled R-squared values, we deemed the full model fit for that cell on that day to be significant. We used Bonferroni correction on the threshold percentile, dividing the p-value by the number of cells in each day, such that the threshold percentile is calculated as 100 - 5/(number of cells for this mouse on this day). See Supplementary Figure 4B,C for counts and proportions of cells with significant predictions, broken down by mouse and day.

#### Fitting trial-average models

All linear models in this study treat each timepoint as an independent observation and regress across all observations in a session. This approach is standard for quantifying the neural encoding of movements, but it conflates encoding for the mean movement with encoding for the variability around the mean. Thus, we compared them to models that predict a neuron’s single-trial activity from trial-averaged movements.

For each predictor separately, we calculated a time-varying average movement during reaching, locked to the onset of reach. This is very similar to calculating a PETH, but where we average across snippets of a kinematic variable, instead of snippets of neural activity. Since reaches are different lengths, the value for each timepoint (relative to the onset of reach) was calculated as the average value across all reaches that were at least that long. Predictor averages were calculated out to the length of the longest reach, including only timepoints that were still part of a reach in the average. We built a new design matrix where for each predictor we replaced each reach with a copy of that predictor’s average. Each reach was replaced with a corresponding number of timepoints of the average: for a reach that is 10 timepoints long, we replaced those 10 timepoints with the first 10 timepoints of the reach-locked average, whereas for a reach that is 20 timepoints long, we replaced those 20 timepoints with the first 20 timepoints of the reach-locked average. Only reach timepoints were used for the models, as for the single-trial models. The time-in-session predictor was treated in the same manner as the behavioral predictors, yielding a predictor that simply increases linearly over the course of a reach. Trial-average models were fit using all behavioral predictors and the time-in-session predictor. Since trial-average models serve as a within-session comparison, we do not perform across-day covariate balancing on trial-average design matrices. All trial-average models were fit and tested for significance using the same cross-validated train-test procedure described for single-trial full models above.

### Comparison of linear encoding models across days

As stated in the main text, for a given cell, when comparing its encoding of trial-by-trial variance across days, we only consider days where the cell has a significant single-trial full model fit. This represents the majority of cell-days recorded in this study (79.7% of all cell-days). Cell-days without significant single-trial full model fits do not show significant tuning for the movement predictors we consider here. Thus, their model fit components do not represent the activity of the cell in a statistically significant way and should not be used to quantify similarity or dissimilarity of encoding.

#### Applying models across days

To apply a model from one day (train day) to data from another day (test day), we weight the predictor vectors from the test day with the coefficients from the train day full model fit and add them together to generate a prediction for test day neural activity (schematized in Fig 6A), for which we calculate an R-squared value. As when fitting the models on their train days, when applying models to test day data, the predictors and neural activity are z-scored before applying, so no constant term is required. We used all cell-days with a significant full single-trial model fit as both train and test days: for a given cell, we applied all significant models to all other days with significant models. To test the applied model R-squared value for significance, we compare to the timepoint shuffle null distribution of the test day, the same distribution used to identify if the test day model itself provides a significant prediction. Forward application refers to when the train day occurs before the test day; backward application refers to when the test day occurs before the train day. Because of our requirement that both test and train days have significant full model predictions, all train and test day pairs have a forward and a backward application. For a given train and test day pair, we interpret the neural encoding as stable across these two days if the application in at least one direction is significant.

When comparing applied model R-squared values to test day trial average R-squared values, we split model applications into two groups based on the relative magnitude of the test day’s single-trial model R-squared and trial-average model R-squared. Note that because of this split, we must report in terms of model applications rather than cell-day pairs. Each cell-day pair constitutes two model applications (forwards and backwards). The two cell-days in a pair might not fall on the same side of this split (one might have a better single-trial fit than trial-average fit, while the other might have a better trial-average fit than single-trial fit), meaning the model applications for this pair will need to be split up.

#### Notes on comparing linear model fits in the presence of correlated predictors

When predictors are sufficiently correlated, as in our study, there is an infinite set of possible coefficient combinations that can produce the same optimal fit in the full models. Even with appropriate regularization, model fits for a given day will be specific to that day’s predictor correlations. If the correlations between predictors change from the training day to the test day, the coefficients derived from the training day may not accurately reflect the predictor relationships on the test day, potentially resulting in a drop in performance on the test day. This drop can occur even if the underlying relationships of interest, e.g. single neuron encoding of individual predictors, remain stable. If predictor correlations are not carefully accounted for, applying fitted models across days may mistakenly suggest that activity-behavior relationships for individual predictors are changing, when in fact, it is the relationships among the predictors that are shifting.

To mitigate this possibility and increase confidence in attributing observed instability to changes in neural encoding rather than changes in behavior, we take the following steps in our across-day analysis: 1) We perform covariate balancing on the movement predictor sets before fitting and comparing linear models. Because this procedure balances the joint distributions of predictors by considering all movement predictors together when matching timepoints, a by-product is partial enforcement of similarity (over days) in the relationships between pairs of predictors. 2) When applying models over days, we do not require model application to be significant in both directions for a pair of cell-days to qualify as having significantly similar encoding. In theory, if there is explanatory power in the predictor set that is maintained across days, this approach allows us to detect that stable encoding even if predictor correlations change meaningfully over days, as illustrated by the following example. On day 1, predictors x_1_ and x_2_ are correlated with each other and also with the predicted variable y in the same way, so model coefficients will give approximately equal weight to x_1_ and x_2_, perhaps with bias to one or the other dependent upon noise. On day 2, predictor x_1_ is still correlated with predicted variable y in the same way, but x_2_ is less correlated with both x_1_ and y or perhaps correlated in a different way, so model coefficients for day 2 will give more weight to x_1_ than x_2_. Applying the model from day 1 to day 2 will not weight x_1_ enough and will weight x_2_ too highly, leading to a poor prediction. However, applying the model from day 2 to day 1 will yield a fit similar to day 1’s own model, since on day 1 the contribution from x_1_ or x_2_ is approximately equivalent. Requiring model application to be significant in both directions requires predictor correlations to be precisely the same on each day in order to detect stable encoding. Not requiring symmetry allows us to detect stable encoding in the face of potentially changing predictor correlations. In practice, we observe that only 7.2% of pairs of cell-days deemed to have significantly stable encoding by this metric are significant in only one direction (328 out of 4586 pairs of cell-days), meaning that the vast majority (92.8%, or 4258 out of 4586 pairs of cell-days) are bidirectionally significant.

Finally, as with all similar studies, we are able to quantify encoding for the movement predictors we included but cannot speak to whether cells significantly encode other unrecorded features of this behavior. There are likely other features of the animals’ behavior that are encoded by a portion of the neural population and are correlated with the variables measured here. As such, in the case of a cell with significant encoding of recorded movement predictors on some days but not all, we are unable to differentiate between the following two situations: 1) that cell encodes the recorded movement features and has lost its relationship to them on some days or 2) that cell encodes different, unrecorded movement features that are correlated with the recorded features on some days but not all. We thus focus our across-day analyses on asking whether cells that maintain significant encoding of our collection of recorded movement variables maintain the nature of their encoding over days.

#### Idealized stable across-day trial-average encoding with linear models

Applying trial-average linear models across days cannot be done with the same rigor as for single-trial linear models, and therefore does not provide a valid test of across-day stability of trial-average encoding. For single-trial models, we use covariate balancing to minimize across-day differences in movement statistics before comparing encoding. This procedure is not possible for trial-average models. The trial-average predictor matrix is constructed from repeated copies of the mean movement trajectory, so its statistics are fixed by definition and cannot be sub-sampled or balanced across days. Consequently, any performance drop observed when applying a trial-average model across days would be fundamentally ambiguous: it could reflect neural encoding drift, or simply small differences in the mean movement trajectory (across any of the 15 predictor dimensions). Instead we assess the stability of trial-average encoding across days using PETHs, which provide a cleaner and more interpretable measure at that level of description.

Although trial-average models applied across days is not meaningful, we can compare across-day single-trial encoding against idealized stable across-day trial-average encoding (Supplementary Figure 5B; see Results section *Linear model encoding profiles of single cells are stable across days*). We have access to the upper bound of across-day trial-average model performance: the test day’s trial-average prediction. Its R-squared value reflects the maximal achievable performance under perfectly stable trial-average encoding and thus serves as a principled benchmark.

## Results

### A dexterous reach-to-grasp task elicits variable yet spatiotemporally coordinated forelimb, digit, and head movements

To investigate the stability of motor cortical encoding of moment-to-moment forelimb and head movements, we trained unrestrained mice on a challenging reach-to-grasp task. Mice were trained to reach for a millet seed placed on a narrow pedestal positioned in one of two or three locations just outside an arena (Fig. 1A,B). Mice self-initiated delivery of a seed by moving a specific required distance away from the reach port (generally 1-2 body lengths away; see Methods for details of seed delivery systems). A critical aspect of the task is positioning the body and head after returning to the reach port, to ensure the pedestal is within a reachable distance and angle. In tandem, the mice must make micro-adjustments to forelimb and digit reaching movements based on body positioning. After returning to the reach port, mice performed a bout of reaches in quick succession during which they either successfully retrieved the seed or knocked it off. Unless otherwise noted, throughout this paper, we treat each reach as a single trial. During a 40 minute daily session, a trained mouse will perform hundreds to thousands of individual reaches (1936 reaches per session +/- 542 [789, 2831]; mean +/- st.dev. [range] across all sessions for all mice, n=25 sessions) and successfully retrieve and consume dozens of seeds (70 seeds +/- 34 [27, 139]; mean +/- st.dev. [range]), achieving an average success rate of 13.5% (13.5% +/- 7.1% [29.6%, 5.6%]; mean +/- st.dev. [range]; Methods; Supplementary Figure 1A).

The pedestal has a diameter similar to that of the seed, making it easy for the seed to be knocked off (Fig. 1B). Because of this and the speed at which the paw approaches the pedestal, the mouse has only one chance to contact the seed—resulting either in a successful grasp or, more likely, knocking the seed off and making it unreachable. Consequently, the task demands precise paw placement and highly coordinated, precisely timed digit control, as only a small range of movements will successfully secure the seed. The narrow pedestal also prevents the mouse from repositioning the seed within its digits after contact. Mice must then maintain their grip while transporting the seed back into the arena, adding to the task’s complexity.

To quantitatively describe digit, paw, and head movements during reaching, we recorded high-speed video on all days and employed markerless keypoint tracking to reconstruct the kinematics of each reach (Fig. 1A,C,D; DeepLabCut; Mathis et al., 2018; Nath et al., 2019). From this keypoint tracking, we extracted the position and velocity of the paw and head centroids and used principal component analysis to estimate proxies for the shape and orientation of the paw and the orientation of the head (Fig. 1C; Methods). The position and velocity of the paw centroid reflect activation of proximal forelimb muscles controlling the elbow and shoulder, while principal components (PCs) of the paw markers reflect activation of distal muscles controlling the wrist and digits. Similarly, the position, velocity, and orientation of the head reflect whole-body posture and potentially activation of shoulder and neck muscles.

Within a 40 minute daily session, trained mice exhibit high reach-to-reach variability in the speed, shape, and duration of digit, paw, and head movements during reaching (Fig. 1E,F,G). This trial-to-trial variability persists even though the mice demonstrate stable success rates during the period of study (Supplementary Figure 1A). As our ultimate goal is to examine the moment-to-moment neural encoding of these movements, we consider this within-session variability to be an advantage of the dataset. This is because it provides good sampling of a wider range of movement space than could be achieved with a more stereotyped behavior. This allows us to more accurately quantify neural encoding within each session and more accurately assess the stability of that encoding across sessions.

### Mice employ a reaching pattern that is consistent across days, despite individual differences in movement specifics

To investigate whether neural coding for reach-to-grasp behavior changes or remains consistent over time, we analyzed data from five consecutive days late in training (starting between day 20 and 22 of training, Methods), a period during which mice demonstrate stable success rates (Supplementary Figure 1A). The within-session variability observed on a given day persists across the five days of study, with variability centered around a similar mean on each day (Fig. 2A). For each mouse, the distributions of all measured kinematic properties remain consistent across all five days. This day-to-day consistency in movement allows us to assess the neural encoding of a similar set of reach, grasp, and carry behaviors on each day, enabling use of this dataset to study encoding stability.

**Figure 2:**
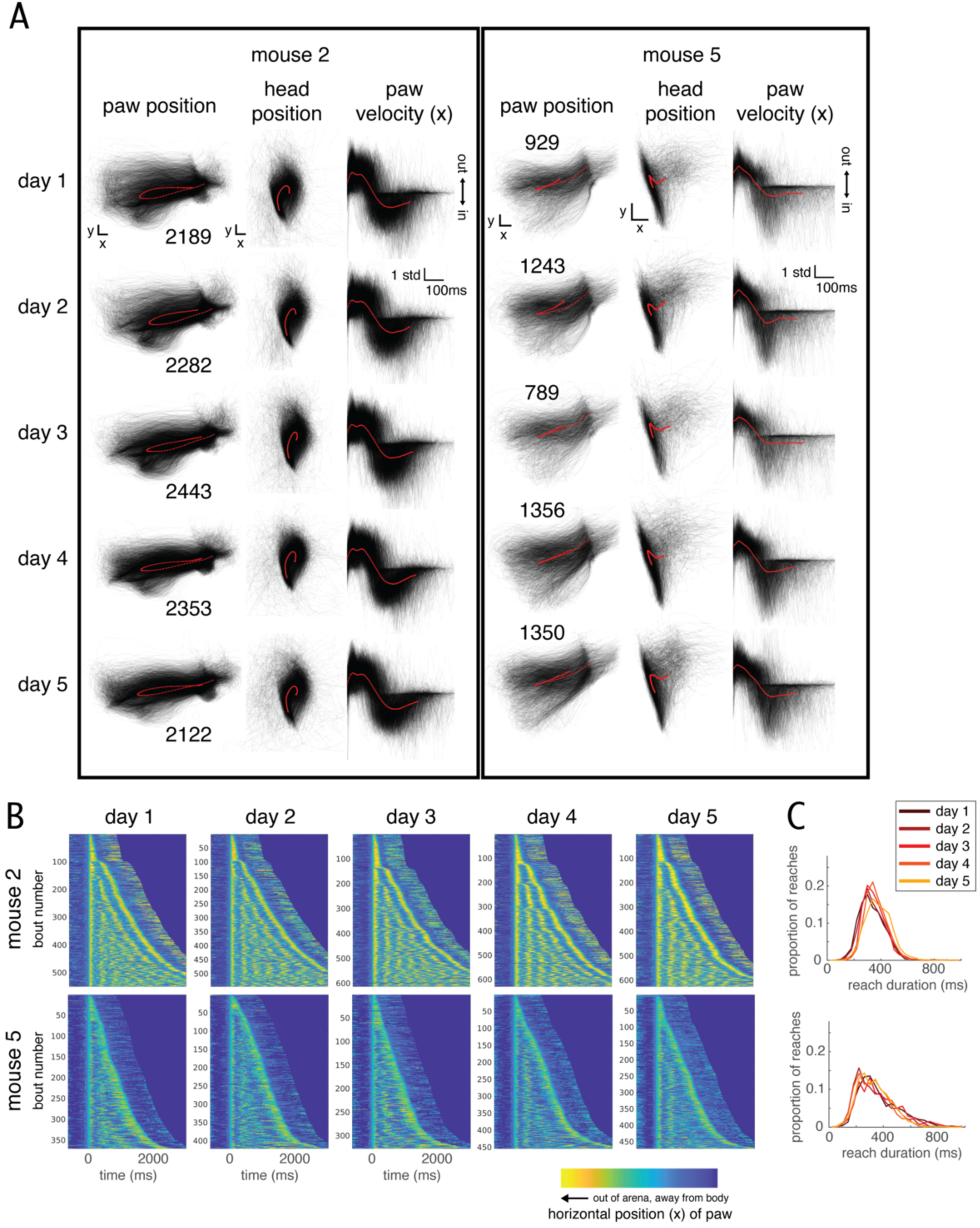
Each animal develops its own unique reaching pattern, yet uses a similar range of movements on each day, enabling study of encoding stability. A) All reaches in each of five daily sessions for two example mice, for 3 of the 14 kinematic variables extracted via keypoint tracking. As in Fig. 1G, individual reaches are plotted in transparent gray, the average (time-locked to reach onset) is plotted on top in red. Numbers next to the paw position plots indicate the number of individual reaches the animal made in that session. B) For the same two example mice as in A (mouse 2 and 5), for all five days: paw position in the x dimension plotted the same way as in Fig. 1E. Colormap corresponds to the position within the frame of the tracking camera, as schematized in Fig 1A. C) For the same two example mice as in A (mouse 2 and 5), for all five days: probability density distributions of time duration of individual reaches, color indicates session within the stability dataset.

The specific digit, paw, and head movements employed during reaching differ across mice, as each mouse developed its own strategy for retrieving the seed (Fig. 2A,B, Supplementary Figure 1B,C). As all of the analyses that follow are performed within-animal, i.e. we do not pool cells across mice, they are not impacted by these observed individual differences in task strategy. Instead, what is vital for these analyses is identification and control of any behavioral differences across days within the same mouse. Ultimately, each analysis yielded results that are consistent across mice, suggesting that the properties of neural coding in question are not strongly dependent on the observed behavioral differences.

### Neurons in forelimb motor cortex (CFA) modulate activity during reaching

We recorded the activity of populations of individual neurons in caudal forelimb area of mouse motor cortex (CFA, or forelimb M1) using an Inscopix miniscope to perform one-photon calcium imaging in freely moving (non-headfixed) mice. We combined the miniscope with a prism to achieve an imaging plane perpendicular to the cortical surface (Andermann et al., 2013; Resendez et al., 2016). This setup enabled simultaneous imaging across the depth of layer 2/3, capturing hundreds of cells in a single field of view (244 cells +/- 50 [138, 347]; mean +/- st.dev. [range], n=25 sessions). By consistently returning to the same field of view each day, we were able to longitudinally track the activity of many of the same cells over multiple days (Fig. 3A,B).

**Figure 3:**
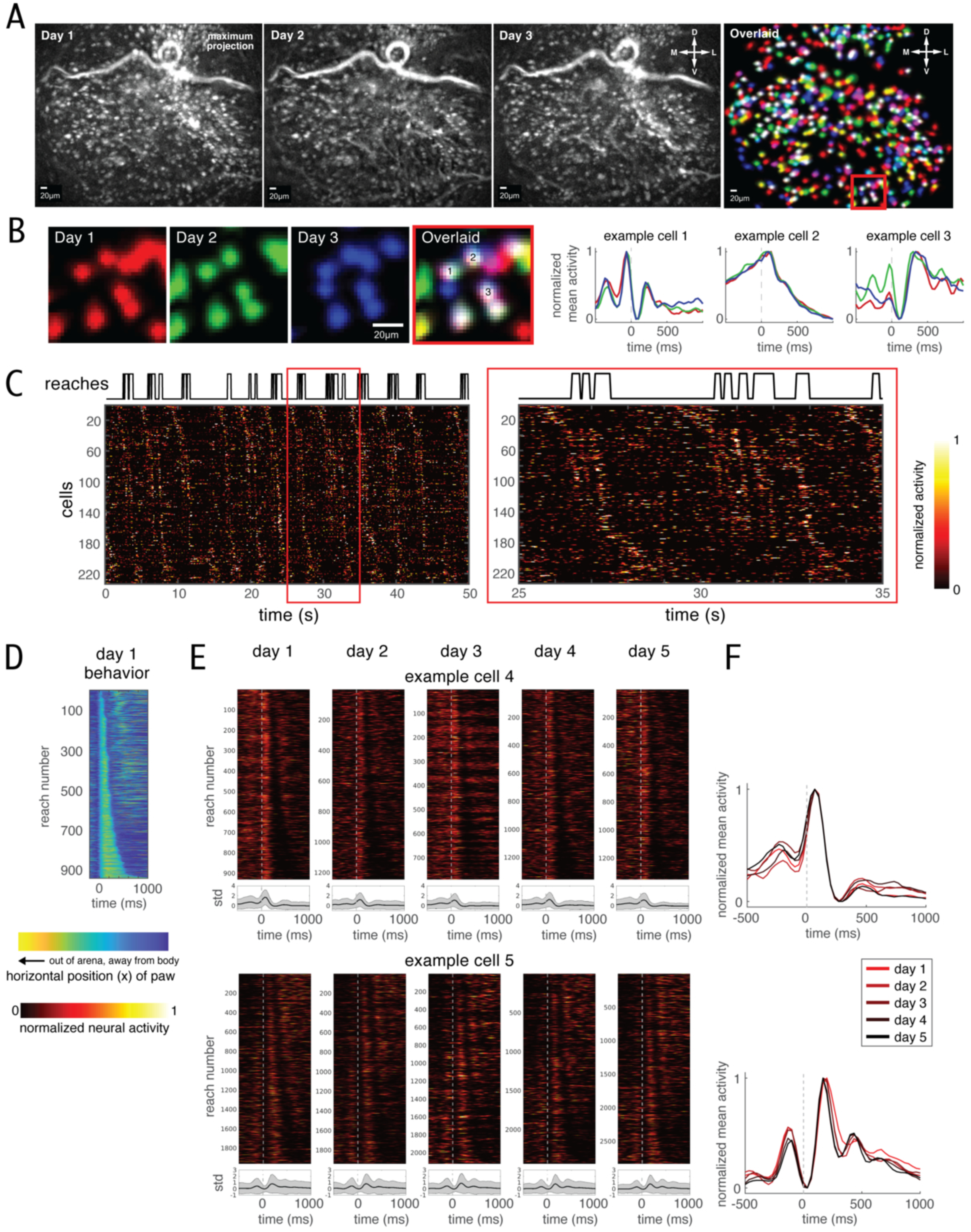
Neurons in forelimb motor cortex (CFA) modulate during reaching. A) Maximum projections of the motion-corrected calcium imaging movie of the same field of view (from mouse 4) on three consecutive days. Bright spots indicate cells; a blood vessel is visible at top. Compass indicates dorsal-ventral and medial-lateral. Far right: overlay of footprints of all individual cells identified on all three days, with day 1 in the red channel, day 2 green, and day 3 blue. White footprints indicate cells found on all three days. B) Tracking of individual neurons across sessions. Left: Zoomed in view of red square outline in A far right, separated by days and also overlaid. Right: peri-event time histograms (PETHs) locked to the onset of reach (see main text) for the three days shown, for three example cells from the zoomed view. PETHs are normalized for each day separately so that minimum value is 0, maximum value is 1. Colors indicate day. t = 0 indicates the time of reach onset. C) Reach modulation is apparent in single-trial neural activity. Top: Binary variable indicating when the mouse was reaching (high indicates reaching, low indicates not reaching). Bottom: Heatmap of inferred spiking activity of all recorded cells from one field of view for 50 seconds of recording. Right inset: Temporal zoom to 10 seconds of recording, corresponding to the red box in the full view. Each cell’s activity has been normalized separately within this window: minimum value = 0, mean + 5*standard deviation = 1. Cells have been sorted according to the time of the peak in their reach-locked PETHs. D) All reaches in an example daily session for an example mouse, copied from Fig. 1E bottom left, for comparing with single-trial neural activity in E and F. As in Fig. 1E, heatmap indicates paw position in the x dimension plotted for all reaches in one daily session for an example mouse (mouse 5). Colormap from Fig. 1A is copied at bottom; yellow colors indicate further outside the arena, blue colors indicate closer to or inside the arena. Reaches are sorted according to reach duration. t = 0 indicates the time of reach onset. E) Individual cells show similar patterns of reach modulation across days. Single-trial inferred spiking activity of two example cells found on all five days, aligned to the start of reach. For each day, top panel shows a heatmap of the single-trial activity. Bottom panel shows the average response across all trials on that day (PETH). Heatmaps are normalized within each day as 0 = mean activity, 1 = 10*standard deviation above the mean. Trials are sorted according to the duration of reaches, as for the kinematics in D: short reaches at the top, long reaches at the bottom. PETHs are plotted as mean +/- st dev, units refer to standard deviations above or below the mean. F) PETHs for example cells in E, from all five days. PETHs are normalized for each day separately so that minimum value is 0, maximum value is 1. Colors indicate day. t = 0 indicates reach onset.

The vast majority of neurons exhibited significant modulation during reaching relative to all other timepoints: 5544 out of 6105 cell-days (90.8%), where each cell-day represents a single day’s observation for a given neuron, which constitutes on average 90.8% of cells in a given session (90.8% +/- 4.0% [82.1%, 96.5%]; mean +/- st.dev. [range]; ANOVA with p = 0.05, with Bonferroni correction for the number of cells in that session (field of view on that day); Methods). Across the recorded population, neurons displayed diverse temporal response profiles, with activity observed before, during, and/or after reaching (Fig. 3C).

Additionally, cells were active during the intervals between reach bouts, which includes other behaviors such as consuming seeds, grooming, and locomotion to trigger the next seed delivery or to position the body at the reach port. In all analysis, each reach was treated as a trial. Neural activity showed notable variability across individual trials (Fig. 3C,E). Not all neurons were active during every reach, and even across reaches when a neuron was active, the magnitude of its activity varied. Despite this variability, neurons demonstrated stable overall patterns of activity across days (Fig. 3E,F), both in the pattern of single-trial activity relative to reach duration and in trial-averaged responses.

### Average neuronal activity patterns locked to reach (PETHs) are stable across days

CFA has recently been shown to encode low-level details of forelimb movement such as muscle activation and kinematics, e.g. joint position and velocity (Omlor et al., 2019; Koh et al., 2025; Grier et al., 2026) suggesting a role in online movement monitoring and possibly control. However, earlier work demonstrated that CFA also encodes high-level task variables such as behavioral category and reach direction (Peters et al., 2014, 2017; Huber et al., 2012; Dombeck et al., 2009; Galiñanes et al., 2018). We thus began our neural encoding analysis by assessing whether CFA encodes a high-level behavioral category signal for a reaching behavior with high trial-to-trial variability. We quantified the average response of each neuron during reaching using peri-event time histograms (PETHs) time-locked to the start of the reach. These PETHs reflect a neural signal corresponding to the behavioral category of reaching, rather than encoding of the details of individual reaches.

The diverse response time courses observed on individual trials (Fig. 3C) were also evident in the trial-averaged responses of individual neurons (Fig. 4A). Despite this diversity, PETHs from the same longitudinally tracked neuron on different days were highly similar, consistent with prior work examining more stereotyped behaviors in rats, mice, and zebra finches (Jensen et al., 2022; Aparicio et al., 2026; Katlowitz et al., 2018). To quantify this across-day similarity, we computed the correlation between each longitudinally tracked neuron’s PETH on pairs of days, using activity from 100ms before to 500ms after reach onset. This window restricted the comparison to timepoints immediately before, during and immediately after reaching (Methods). For each mouse, we examined the population distribution of across-day correlation values as a function of the interval between days (Fig. 4B,C; Methods).

**Figure 4:**
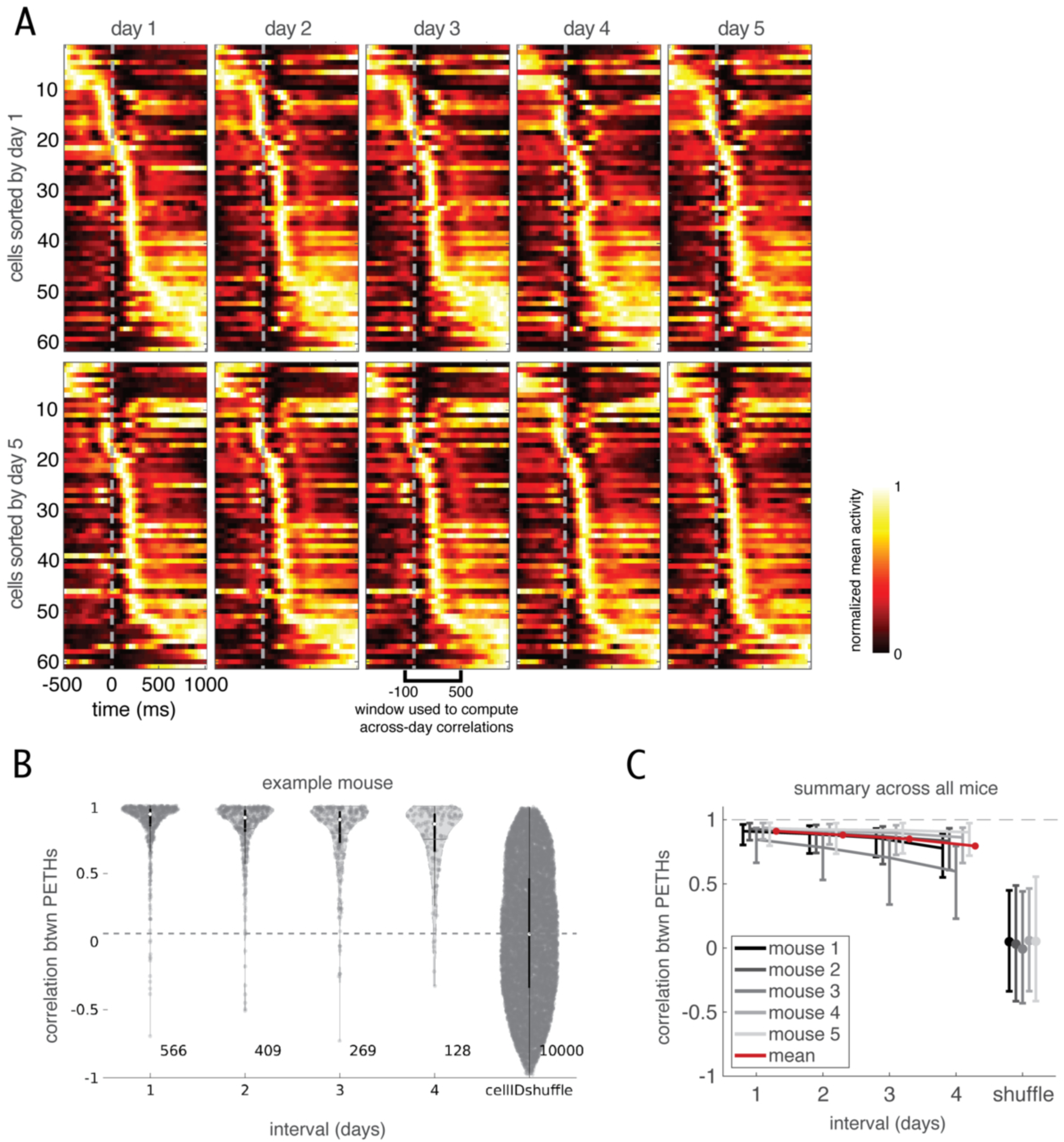
Average neuronal activity patterns (PETHs) are stable across days. A) Normalized peri-event time histograms (PETHs) of neurons found on all five days for an example mouse (mouse 4). Cells are sorted according to the time of their peak activity on either day 1 (top row) or day 5 (bottom row). Each cell’s PETH has been normalized separately within each day within the plotted window: minimum value = 0, maximum value = 1. t = 0 indicates the time of reach onset. B) Correlation between PETHs, as a function of the interval between the days, for all cells found on more than one day for an example mouse (mouse 4). Each datapoint is an observation of the same cell on two different days. Numbers to the bottom right of the violins indicate the number of observations included in that violin. For each violin, the white dot indicates the median of the distribution and the black box indicates interquartile range. Cell ID shuffle indicates the distribution of correlations between PETHs of two cells randomly chosen from any day (excluding any cell pairs that are registered as the same cell; see Methods). Gray dashed line indicates the median of this shuffle. C) Correlation between PETHs for all mice. Datapoint is median, error bars are interquartile range across cells for that interval for that mouse. Grayscale indicates mouse. Shuffle as in B.

For all intervals, the distributions were strongly shifted toward positive correlations, indicating that during reaching neural activity patterns are generally preserved across days. The average median correlation was 0.91 for 1-day intervals, 0.88 for 2-day intervals, 0.85 for 3-day intervals, and 0.79 for 4-day intervals (average computed across n=5 mice). To contextualize the magnitude of this similarity, we generated a control distribution by computing the correlations between PETHs for different neurons rather than from the same longitudinally tracked neuron (Fig. 4B; Methods). The medians of the observed correlation values for each interval was far from the median of the control distribution, which was close to zero (Fig. 4B,C). Thus, across the population, an individual neuron’s activity pattern was more similar to its own activity across days than to the activity patterns of other neurons. This remained true even across a four-day interval, indicating that the trial averaged spatiotemporal pattern of population activity during and around reaching is largely conserved within the period of study.

Despite this overall stability, across-day coding similarity decreased modestly as the interval between days increased (0.12 +/- 0.08 [0.02, 0.25]; mean +/- st.dev. [range]; absolute difference between medians computed within mice, then averaged across mice, n=5 mice). Consistent with this gradual decrease, the correlation distributions for the 1-day interval and the 4-day interval are statistically differentiable for 4 out of the 5 mice (p < 0.05, Kolmogorov-Smirnov (KS) test performed within animal).

Importantly, this reduction in coding similarity was not evenly distributed across the population. A large portion of cells maintained high across-day similarity, with across-day correlations that remained near one even over longer intervals. However, at all intervals considered, a portion of cells showed lower across-day similarity. The correlation distributions therefore broadened rather than simply shifting uniformly toward lower values. This pattern suggests heterogeneity in apparent drift rates: some neurons maintained highly stable reach-related activity patterns, whereas others changed more substantially across days. Longer-duration longitudinal recordings will be required to determine whether the persistence of high correlations for a portion of the population reflects a ceiling effect of the correlation metric, in which case all neurons would eventually show reduced coding similarity, or instead indicates the existence of a subset of neurons with little or no measurable drift over behaviorally relevant timescales. Regarding the subset of neurons that show decreased correlations, longer recording durations will be required to determine whether the drop in PETH correlations over a four-day interval reflects a slow but steady drift that would continue to accumulate or whether coding similarity would eventually stabilize. A gradual reduction of coding similarity over short intervals followed by a stabilization at longer intervals was observed in rat motor cortex and dorso-lateral striatum (Jensen et al., 2022). Such a pattern may reflect the influence of processes that fluctuate around a long-term stable mean with an autocorrelation timescale greater than a few days, e.g. physiological factors such as hormonal fluctuations or uncontrolled environmental factors. Finally, we note that trial averaging is not ideally suited to a behavior with high trial-to-trial variability, motivating the single-trial encoding analyses described in the next section. Indeed, given the within-session behavioral variability, the reliability of the trial-averaged signal is notable.

The reach-locked PETH correlation analysis is performed treating each reach as a single trial, but we can also consider each bout as a single trial. To assess the stability of neural encoding of a bout, we perform the same correlation analysis with PETHs locked to the onset of a bout, defined as the onset of the first reach in the bout (Supplementary Figure 3A-C; Methods). We found lower levels of stability for bout-locked PETHs than for reach-locked PETHs. The average median correlation was 0.85 for 1-day intervals, 0.82 for 2-day intervals, 0.77 for 3-day intervals, and 0.74 for 4-day intervals (average computed across n=5 mice).

Across-day coding similarity decreased as the interval between days increased (0.11 +/- 0.07 [0.02, 0.21]; mean +/- st.dev. [range]; absolute difference between medians computed within mice, then averaged across mice, n=5 mice). The across-bout variability in movement is inherently even greater than the across-reach variability, owing to differences in number of reaches, reach timing within the bout, and related structure (illustrated for one kinematic dimension in Fig. 1E, Fig. 2B, and Supplementary Figure 1C). Because of this, we cannot determine whether the lower correlation for bout-locked PETHs reflects less stable neural encoding or greater intrinsic variance in the constituent movements being averaged over.

### Trial-by-trial variance in forelimb, digit, and head movements is captured in single-trial neuronal activity in CFA M1 during a dexterous reach-to-grasp task

Paw and head movements during this reach-to-grasp task exhibit trial-to-trial variability across multiple kinematic dimensions including speed, duration, timing of wrist pronation, and timing of digits opening and closing (Fig. 1 & 2). While PETHs quantify a neuron’s response for the average movement, they do not reveal whether a neuron encodes the significant trial-by-trial variance in this behavior, i.e. whether the neuron’s activity changes when a given movement variable changes. Single-trial variance is statistically necessary for quantifying the neural encoding of these continuous variables because it allows us to assess that relationship over a wider range of movement values than the narrow range afforded by more stereotyped behaviors. To assess the neural encoding of the moment-to-moment details of reach-to-grasp kinematics, we constructed families of linear models to predict the activity of individual neurons during reaching based on the extensive set of kinematic variables we extracted using keypoint tracking (schematized in Fig. 5A; Methods). Here we use an encoding approach, predicting a cell’s activity from a set of movement variables), rather than a decoding approach which predicts movement variables from population activity. This choice allows us to assess whether individual cells maintain or change their movement encoding across days in a more straightforward manner. Motor cortical encoding of moment-to-moment forelimb kinematics has been demonstrated in head-fixed mice (Koh et al., 2025; Grier et al., 2026). We extend this work by characterizing forelimb encoding in freely-moving animals, which enables direct comparison with the encoding of unrestrained head movements.

**Figure 5:**
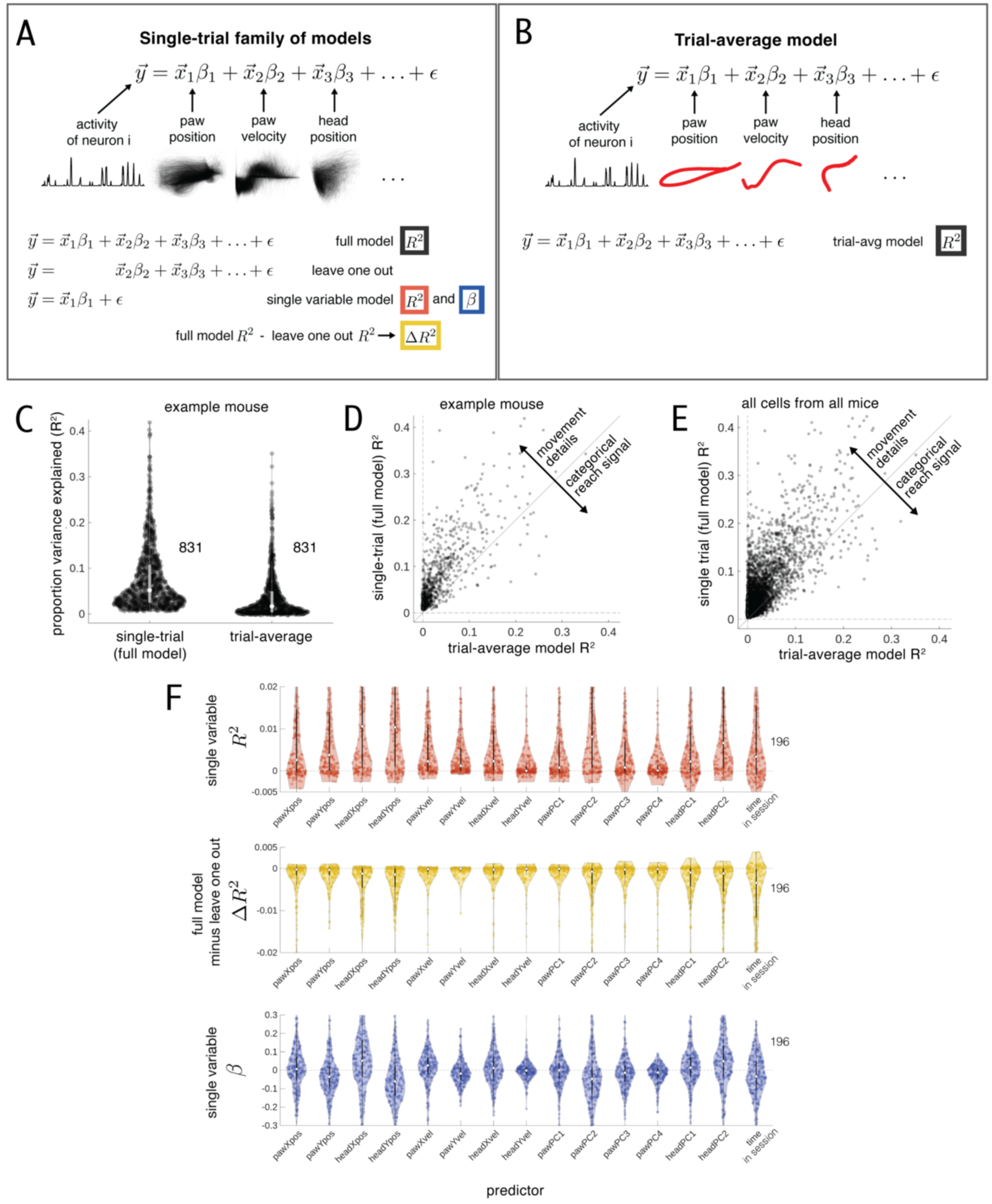
Single-trial variance in forelimb, digit, and head movements allows for accurate modeling of single-trial CFA M1 neuronal activity. A) Schematic of single-trial linear model families built to predict the activity of individual neurons from paw, digit, and head movements during reaching. The full single-trial model contains all extracted kinematic variables and their observed values across trials. For each predictor, we build a leave-one-out model using all predictors except the one in question, and a single variable model using only that predictor. The difference in R-squared value between the full model and the leave one out model indicates the unique variance explained by that predictor. B) Schematic of the trial-average linear model, built to assess how much neural variance is explained by the trial-averaged movements. In this model, each trial of the kinematics is replaced by the average timecourse for that kinematic predictor, averaged across reaches, locked to reach onset (see main text and Methods). C) Proportion of variance explained (R-squared) by the full single-trial model or the trial-average model for all cell-days with a significant full single-trial model prediction, for an example mouse (mouse 5). Numbers to the right of each violin indicate the number of cell-days in the distribution. Each datapoint corresponds to a cell on one day (cell-day). Cells registered across days have multiple dots if they have significant full single-trial model predictions on multiple days. For each violin, the white dot indicates the median of the distribution and the gray box indicates interquartile range. D) Same data as in C, but with the full model and trial-average model R-squared values plotted against each other for each cell-day. Cells that fall above the unity line show better predictions from the single-trial model than from the trial-average model, indicating that trial-averaged movements cannot fully explain their encoding of single-trial movement details. Cells that fall below the unity line are better explained by trial-averaged movements than by single-trial movement details, indicating that their activity is best explained as a categorical reach signal that is similar across all reaches and does reflect the variability of individual reaches. E) The majority of neurons show improved predictions from the single-trial model relative to the trial-average model, indicating that they encode single-trial movement details. Same as D, but for all cell-days with significant full model predictions from all mice. F) Model features from the single-trial family of models for all cells from one example session from one mouse, for each predictor separately (all kinematic variables and time within the session; see main text and Methods for additional details of predictor sets and variance explained metrics). Each kinematic predictor contributes predictive power (non-zero variance explained) for some portion of neurons, and single neurons show a range of linear relationships with movement variables. Within a violin, each datapoint corresponds to a cell, n = 196 cells with significant full single-trial model fits for this session; each cell has exactly one datapoint in every violin. For each violin, the white dot indicates the median of the distribution and the gray box indicates interquartile range. Top: variance explained (R-squared) for single variable model fits. Middle: same as top but for the delta-R-squared, indicating the unique variance contributed by each predictor. Bottom: same as top but for the coefficients from single variable model fits, reflecting the linear mapping from each predictor to neural activity independent of the predictor’s correlation with other predictors.

Notably, neural activity could encode the mean movement without encoding the trial-by-trial variability. This would suggest that motor cortex carries a general ’reach’ signal—indicating whether the mouse is performing the reaching behavior or not—without capturing the moment-to-moment details of the movements involved. While individual neurons exhibit significant trial-by-trial variance around their average responses (Fig. 3D), it is possible that that variance does not co-vary with the variance observed in movement.

Additionally, even if movement details are indeed encoded in motor cortical activity on a given day, it is also possible that the nature of that encoding is not the same across days, indicating instability (drift).

Before fitting the models, we applied a covariate balancing subsampling procedure to the kinematic predictor sets for all five days, separately for each mouse, to ensure that we are comparing encoding of similar movements across days (Supplementary Figure 2; Methods). This step is necessary for later analysis of encoding stability; without it, we would not be able to differentiate between true drift and changes in encoding caused by changes in behavior. For all mice, covariate balancing on the kinematic predictor joint distributions increased the within-mouse, across-day similarity of individual predictor distributions for the majority of predictors (560 out of 710 individual day-pair comparisons (78.9%); 60 out of 75 predictors when Wasserstein distance is averaged across day-pair comparisons but not across mice (80.0%); Supplementary Figure 2; Methods). Only timepoints during reaching are used in the linear model analysis, meaning that timepoints occurring between individual reaches, while the animal is away from the reach port, or while the animal consumes seeds are not included.

The majority of neurons on any given day have a modest but statistically significant amount of variance explained when predicted by the entire set of kinematic predictors, as compared to a timepoint-shuffled control (full single-trial model schematized in Fig. 5A). Specifically, 4871 out of 6105 cell-days (79.8%) showed significant variance explained by the full single-trial model, where each cell-day represents a single day’s observation for a given neuron (p = 0.05, with Bonferroni correction for the number of cells in that session [field of view on that day]; Methods). On average, this comprises 79.6% of neurons in a given session (79.6% +/- 19.5% [39.1%, 96.0%]; mean +/- st.dev. [range], n=25 sessions; see Supplementary Figure 4B,C for counts and proportions per session and mouse). For the 4871 cell-days with significant variance explained by the full single-trial model, the median R-squared value is 0.031; on average, the median R-squared value across all cell-days in a session is 0.032 (0.032 +/- 0.011 [0.017, 0.060]; mean +/- st.dev. [range]).

We have already observed that individual cells exhibit time-varying average responses (PETHs in Fig. 4), indicating encoding of the trial-averaged movement. To contextualize the magnitude of encoding of single-trial variance relative to encoding of the trial average, we compare performance of the full single-trial model to that of a trial-average model, in which we predict the cell’s single-trial activity from trial-averaged movements (schematized in Fig. 5B). In the predictor matrix of this trial-average model, we replace the timepoints of each reach with the reach-locked average, for each kinematic predictor separately (Methods). The majority of recorded neurons were significantly predicted by a trial-average model (4312 out of 6105 cell-days, or 70.6%; Methods), corresponding to on average 70.4% of neurons in a given session (70.4% +/- 15.1% [31.7%, 93.6%]; mean +/- st.dev. [range]). Almost all of these were also significantly predicted by the full single-trial model (3927 out of 4312 cell-days, or 91.1%). Few neurons were significantly predicted by only the trial-average model, indicating they encode only a general reach signal (385 out of 6105 cell-days, or 6.3%), or by only the full single-trial model, indicating they encode only moment-to-moment details (944 out of 6105 cell-days, or 15.5%). For the 4312 cell-days with significant variance explained by the trial-average model, the median R-squared value is 0.014; on average, the median R-squared value across all cell-days in a session is 0.015 (0.015 +/- 0.007 [0.005, 0.030]; mean +/- st.dev. [range]). These and all other linear model results are qualitatively similar when neural activity is predicted from kinematic predictors only, excluding the time-in-session predictor (Supplementary Figure 5C-J and Table S1).

Of the neurons significantly predicted by both the full single-trial model and the trial-average model, we found that the majority were better modeled by the full single-trial model than by the trial-average model (Fig. 5C-E; 3371 out of 3927 (85.8%) cell-days, 85.0% +/- 7.6% of neurons in a given session [59.7%, 94.5%]; mean +/- st.dev. [range]), indicating that for many cells single-trial variance is more strongly encoded than the average reach. Together, these results indicate that mouse motor cortex encodes not only a general “reaching” signal but also moment-to-moment details of movement. For the remainder of our analysis, we focus on the subset of neurons with significant full single-trial model predictions.

Next we consider the contribution of individual predictors. Due to the nature of the task and the physics of the mice’s bodies, many kinematic variables are correlated (Supplementary Figure 4A). Thus, the coefficients in our full single-trial model should not be interpreted as the maximum contribution of each individual kinematic predictor (Stevenson et al., 2018). To assess the contribution of individual predictors to explaining a neuron’s activity, we compute two metrics (schematized in Fig. 5A): 1) Single-variable

R-squared values: For each predictor, we fit a model using only that predictor to estimate the total variance it explains. 2) Delta-R-squared values: Starting with the full single-trial model, we create a leave-one-out model for each predictor by excluding it and refitting the model. The resulting drop in R-squared indicates the unique variance that predictor contributes. Additionally, to capture how a neuron’s activity is modulated relative to variations in each movement predictor, we also consider coefficients (betas) from the single variable models, which serve as a form of tuning curve.

Many predictors exhibit non-zero single-variable R-squared values and delta-R-squared values (Fig. 5F top, middle), indicating that these movement variables are encoded by single cells in motor cortex.

Additionally, distributions of single variable model coefficients are dispersed away from zero (Fig. 5F bottom). Notably, despite partial correlation with paw-related predictors (Supplementary Figure 4A), several head-related predictors have non-zero delta-R-squared values for many neurons, indicating that they contribute unique explanatory power. Thus, the moment-to-moment details of the reaching movements encoded in caudal forelimb area of motor cortex include not only the position of the paw but its shape and orientation (posture of the digits and wrist) and the position and orientation of the head.

### Linear model encoding profiles of single cells are stable across days

To assess whether the encoding of trial-by-trial variability is consistent over time, we tested whether the linear models generalized to days other than the ones they were trained on (Fig. 6A; Methods). Specifically, we applied single-trial full models forward and backwards, restricting our analysis to cell-days that showed statistically significant movement encoding—that is, for a given cell we included a day in this analysis if it had a significant single-trial full model prediction (trained and tested on that day; Methods). For cells with significant fits on some days but not others, we only compared days with significant fits, which constitutes 79.8% of all cell-days (see previous section). Model performance declined only slightly as the interval between training and testing days increased (Fig. 6B). Of the 4,961 cell-day pairs examined across all intervals, 4,596 (92.6%) showed significantly similar encoding, defined as having at least one significant forward or backward model application (Methods), with significance assessed relative to the time-shuffled control for the test day. The vast majority of these—4,283 pairs (93.2% of significant pairs, or 86.3% of all pairs)—exhibited significant encoding in both directions. The average proportion of cell-day pairs that show significantly similar encoding in at least one direction is 94% for 1-day intervals, 91% for 2-day intervals, 90% for 3-day intervals, and 88% for 4-day intervals (Fig 6D; average computed across n=5 mice). The proportion of cell-day pairs that show significantly similar encoding is greater than 80% for all intervals for all mice within this study.

**Figure 6:**
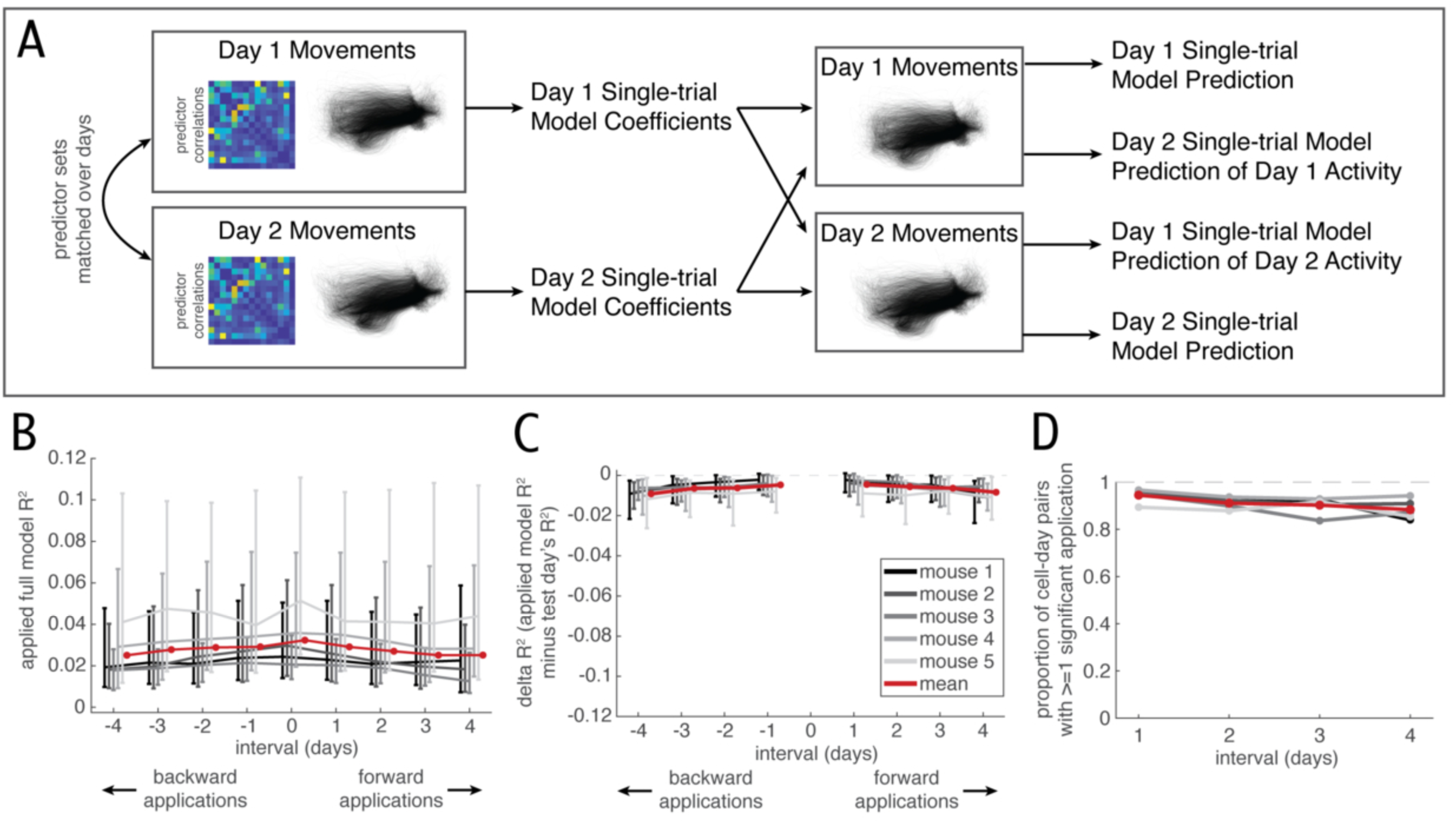
Linear encoding models stably predict single neuron activity across days. A) Schematic of applying linear models across days to assess the stability of single-trial encoding: The full single-trial linear model built for a given cell on a given day is applied to other days for that cell. Kinematic variable predictor sets are matched before fitting models (see main text and Methods). Only cell-days with significant single-trial full model predictions are included in across-day linear model analysis. B) Resulting R-squared value when a model fitted on one day is applied to another day, for all intervals between train and test days. Datapoint is median, error bars are interquartile range across train-test day pairs for that interval for that mouse. Grayscale indicates mice, as in C. Forward applications indicate the train day comes before the test day; backward applications indicate the train day comes after the test day. Distributions include all cell-day pairs where both the train and test day have significant (full) single-trial model predictions. C) Performance of applied models relative to a test day’s own model. Difference between R-squared from an applied model and the R-squared of the test day’s own fitted model. Datapoint is median, error bars are interquartile range across train-test day pairs for that interval for that mouse. Forward and backward applications as in B. Distributions include all cell-day pairs where both the train and test day have significant (full) single-trial model predictions, the same as in Fig. 6B. D) The majority of applied models yield statistically significant predictions. Proportion of cell-day pairs with significant model application in at least one direction, as a function of interval between the cell-days. Grayscale indicates mice, as in C. Applied models are compared to test day’s own timepoint shuffle control to determine significance (see Methods).

Having significantly similar encoding across an interval does not imply that a neuron’s encoding of movement is perfectly invariant. To compare neural encoding in a more nuanced way, we quantify the drop in model performance when applied over days. We computed a delta-R-squared value as the difference between the R-squared value for the test day’s own single-trial model and the R-squared value for the applied single-trial model when applied to test day data (Fig. 6C). The R-squared value for the test day’s own single-trial model reflects the best prediction that could be achieved with test day single-trial data, so the magnitude of these delta-R-squareds reflects how much single-trial encoding changed.

As the interval between training and testing days increased, the magnitude of these delta-R-squared values increased modestly (0.004 +/- 0.002 [0.002, 0.007]; mean +/- st.dev. [range]; absolute difference between medians for 1-day and 4-day intervals, computed within mice, then averaged across mice, n=5 mice). We find that the distributions of delta-R-squareds for 1-day and 4-day intervals are statistically distinguishable for 4 out of the 5 mice (p = 0.05, Kolmogorov-Smirnov (KS) test performed within animal), indicating that, at the population level, there is a modest but quantifiable change in single-trial encoding over the five days in our dataset. Similar to the PETH correlations, we observe that the distributions of delta-R-squared values broaden rather than shifting uniformly downwards, suggesting that drift rates are heterogeneous across neurons. Again, although we observe that a large fraction of cell-day pairs show significantly similar encoding within the window of this study, longer recording durations are required to differentiate between two possibilities: one, that all neurons have non-negligible drift rates and would eventually show reduced coding similarity as within-day intervals increased, and two, a portion of neurons have low to negligible drift rates and would maintain significantly similar encoding over long timescales. Additionally, it remains unclear whether the drop in delta-R-squared values observed over a four-day interval reflects a slow but steady drift that would continue to accumulate, or whether coding similarity levels would stabilize at longer intervals, as observed for PETH correlations in Jensen et al. (2022).

To further contextualize the magnitude of the observed drop in model performance, we compared the level of drift observed in single-trial encoding to idealized stable trial-average encoding. Specifically, we compared each applied single-trial model’s performance to the test day’s trial-average model performance. The test day’s trial-average model R-squared value reflects the best prediction that could possibly be achieved from applying other days’ trial-average models to the test day, and hence represents the idealized case of perfectly stable trial-average encoding (see Methods subsection *Idealized stable trial-average encoding with linear models* for an explanation of why we do not apply trial-average models over days). This value typically exceeds the test day’s timepoint shuffle significance threshold (Supplementary Figure 5A; higher for 3295 out of 3927 cell-days with both significant single-trial encoding and significant trial-average encoding, or 83.9%).

This analysis is restricted to model applications for which the train and test days both have significant single-trial encoding and significant trial-average encoding, meaning those involving the 3927 out of the 4871 cell-days with significant single-trial model fits (80.6%), corresponding to 3833 out of the 4961 *pairs* of cell-days originally examined with across-day linear model application (77.3%). Since each pair of cell-days corresponds to two model applications (one forward and one backward) this corresponds to 7666 model applications. For clarity, we divide these model applications into two groups: those with test days with greater trial-average encoding than single-trial encoding (1201 out of 7666 model applications, or 15.7%), and those with test days with greater single-trial encoding than trial-average encoding (6465 out of 7666 model applications, or 84.3%). This analysis is not well-suited for test days with greater trial-average encoding, as the applied single-trial model obligatorily will perform worse than the trial-average model because it can never do better than the test day’s own single-trial model. Hence, we focus the analysis on the test days with greater single-trial encoding, which constitutes the majority of model applications. We find that 5392 out of these 6465 model applications (83.4%) have an applied model R-squared value that is higher than the test day’s trial-average model R-squared value (Supplementary Figure 5B). The average proportion is 86% for 1-day intervals, 81% for 2-day intervals, 79% for 3-day intervals, 75% for 4-day intervals (average computed across n=5 mice). In other words, for most neurons that encode single-trial variance more strongly than trial-averaged movement, across-day single-trial encoding remains higher—even after four days—than idealized stable trial-average encoding. With this benchmark in place, we interpret the observed drift in single-trial encoding as modest. These and all other linear model results are qualitatively similar when neural activity is predicted from kinematic predictors only, excluding the time-in-session predictor (Supplementary Figure 5C-J and Table S1).

## Discussion

We investigated how motor cortex encodes movements of the paw, digits, and head as mice performed a self-paced, skilled reach-to-grasp task, and we then assessed how stable this encoding is over days. This challenging but flexible motor skill was characterized by many repetitions within a day and substantial reach-to-reach variability—making the skill more reflective of real-world motor behavior. We find that during freely-moving reach-to-grasp, neurons in the mouse motor cortex encode both a general reaching signal and the detailed movements of the head, forelimb, and digits. Notably, both types of encoding remain remarkably stable over days for the majority of individual neurons, providing a reliable neural basis for producing flexible, skilled movements that are both consistent and adaptable.

These findings extend our understanding of how the motor nervous system maintains the ability to produce learned, goal-directed movements. While previous studies in rodents and songbirds have demonstrated that the brain can maintain stable neural activity patterns associated with stereotyped behaviors, i.e. a motor tape (Jensen et al., 2022; Aparicio et al., 2026; Katlowitz et al., 2018), here we demonstrate that it also maintains stable representations of a more flexible behavior. Additionally, while previous work in rhesus macaque has demonstrated stable encoding of single-trial movement details in motor cortex (Gallego et al., 2020), it remains to be seen if the low-dimensional projections of population activity used in that study are the result of stable single cell activity. The results presented here establish two key points: first, that rodent motor cortex shows stable encoding not just of trial averages, but also of single-trial variability of movement details, similar to observations in non-human-primate motor cortex; and second, this stability is maintained at the level of single cells. Such stable encoding of movement details is vital for the production of flexible skilled movements like the one studied here. To enable adaptive control, both online (e.g. within a reach) and offline (e.g. adjustments from one reach to the next), the nervous system must process moment-to-moment details of the constituent movements of complex motor skills. Our findings indicate that the motor nervous system supports flexible motor skills through stable representations of these details, providing a foundation for robust but adaptive performance in dynamic contexts over extended periods of time.

While we observe high across-day coding similarity, the recorded population was not uniformly stable. Notably, the magnitude of observed drift is not uniform across cells. The distributions of PETH correlations and delta-R-squared values broadened rather than simply shifting uniformly toward lower stability, suggesting heterogeneity in apparent drift rates. Longer-duration longitudinal recordings will be needed to determine whether this pattern reflects a ceiling effect, such that all neurons would eventually show reduced across-day similarity, or instead indicates the presence of a subset of neurons with little or no measurable drift over behaviorally relevant timescales. Further work is necessary to determine whether and how the modest and heterogenously distributed drift we observe here would impact decoding of behavioral category and movement details.

In sensory and association brain areas, the neural encoding of stimulus and task variables often shows session-to-session variability, a phenomenon termed representational drift (reviewed in Micou and O’Leary, 2023; Driscoll et al., 2022; Chambers and Rumpel, 2017; Clopath et al., 2017; Lütcke et al., 2013). In contrast, motor areas have been shown to exhibit more stable encoding in a range of species and behaviors (Jensen et al., 2022; Aparicio et al., 2026; Katlowitz et al., 2018; Dhawale et al., 2017; Gallego et al., 2020; Chestek et al., 2007; Fraser and Schwartz, 2012; Flint et al., 2016; but see Rokni et al., 2007 and Liberti et al., 2016). The results presented here corroborate previous findings of stability of motor encoding, and extend this body of work to more flexible and dexterous behaviors. At present, more quantitative comparison of drift magnitude across brain regions remains challenging because reported stability metrics depend strongly on the external variables modeled, the encoding framework used, whether behavioral state has been matched across sessions, the neuronal inclusion criteria, and the controls used to define stability or drift. Further work is necessary both to establish standard approaches for the longitudinal study of encoding and also to determine how the relative stability of motor regions is compatible with the apparent instability observed in sensory and association regions.

Quantifying neural encoding during naturalistic behaviors with high trial-to-trial variability poses significant challenges, which we overcame through a series of carefully designed approaches. First, we used methods to quantify neural encoding that do not assume stereotypy in movements across trials. This analytical flexibility allowed us to account for the inherent variability of naturalistic behaviors. Second, we designed the task to balance behavioral variability with sufficient trial counts. Key elements of the task, such as requiring the mice to reposition themselves after each bout of reaches, randomizing seed positions across multiple locations, and omitting penalties for failed attempts, encouraged both high trial counts and trial-to-trial variability while discouraging the development of stereotyped, automatized movement patterns. The task’s inherent difficulty likely further contributed to the diversity in movements. Third, we captured these movements in fine-grained detail using high-speed, high-resolution cameras, allowing us to precisely quantify trial-to-trial variability above the noise threshold. High-resolution behavioral data is crucial to accurately identify variations in movement encoding. Finally, to ensure that our stability analysis was not confounded by behavioral shifts over days, we implemented a covariate balancing procedure before fitting linear models.

This approach allowed us to quantify encoding within a consistent region of the high-dimensional kinematic parameter set each day, effectively removing neural changes resulting from behavioral variability. Together, these strategies enabled us to rigorously estimate and compare neural encoding across days in a statistically robust manner. Our findings not only highlight the stability of motor cortical encoding during complex, variable behaviors but also underscore the value of incorporating detailed behavioral analyses into longitudinal neural encoding studies (Sadeh and Clopath, 2022; Musall, Kaufman et al., 2019).

We used single-trial, single-neuron encoding models to investigate whether the trial-to-trial variability in movements is encoded in motor cortex and whether this encoding remains stable over days. While more complex encoding models could likely explain additional neuronal variance, they often come at the cost of interpretability. In contrast, our linear approach allowed us to directly link the activity of individual neurons to a carefully curated set of 14 kinematic variables and allowed us to compare the strength of encoding of the movements of specific body parts in a straightforward way.

The finding that motor cortex encodes single-trial variability in reach-to-grasp movements suggests a role in the monitoring and possibly online control of this sensory-driven behavior, likely operating as part of broader sensorimotor loops involving other cortical and subcortical regions such as somatosensory cortex, thalamus, brainstem, and cerebellum (Yamawaki et al., 2021; Sauerbrei et al., 2020; Wagner et al., 2019; Becker and Person, 2019). Further work is needed to determine whether this encoding reflects movement control signals for the ongoing reach, as observed in cerebellum (Becker and Person, 2019) and/or corollary discharge, useful for longer-term processes such as reach-to-reach adjustment (Becker, Calame, et al., 2020; Dhawale et al., 2019) or continual learning as task demands change (Dhawale et al., 2019; Kawai et al., 2015; Hwang et al., 2019).

Our linear encoding model analysis showed that principal components of paw markers, which reflect wrist and digit states, contributed explanatory power comparable to paw position, which reflects the state of proximal joints such as shoulder and elbow. This result implicates CFA not just in precision reaching but also in grasping, contrary to previous results (Wang et al., 2017). Interestingly, position and orientation of the head also contributed explanatory power comparable to paw- and digit-related predictors. This could reflect two mutually compatible explanations: first, head-related variables may be correlated with omitted kinematic variables that are more directly linked to neural activity (Stevenson et al., 2018). Second, head position and orientation may be used in computations in motor cortex, providing crucial information about body positioning relative to the reach port and pedestal. This latter explanation implies a role for egocentric planning signals in guiding the paw to the pedestal, as observed in encoding of whole-body posture in rats (Mimica et al., 2018). These findings highlight the importance of examining the encoding of multiple body parts simultaneously in a given brain area, rather than limiting the focus to movements traditionally associated with that area. A comprehensive understanding of whole-body movement generation requires this broader perspective.

To uncover how the brain produces the flexible, context-dependent, and coordinated movements essential for daily human activities, it is crucial to study comparable behaviors in animal models. The freely-moving reach-to-grasp behavior analyzed here exemplifies such complexity, combining whole-body posture with precise forelimb and digit control. This task closely parallels primate reaching and grasping behaviors (Sacrey et al., 2009; Whishaw et al., 1992; Guo et al., 2015; Becker, Calame, et al., 2020; Whishaw et al., 2017) and serves as a powerful model for investigating the neural mechanisms underlying these intricate actions.

## Acknowledgements

This work was supported by National Science Foundation Graduate Research Fellowship Program (NSF GRFP) fellowship DGE-1746045 (E.A.d.); and National Institutes of Health grant R01-NS094184 (J.N.M). Any opinions, findings, and conclusions or recommendations expressed in this material are those of the author(s) and do not necessarily reflect the views of the National Science Foundation. The authors would like to thank Gabriella Wheeler Fox, Zach Morrissey, Siwei Wang, Harold Rockwell, Sunnie Hong, Steven A. Redford, Stephanie E. Palmer, Nicholas G. Hatsopoulos, Matthew T. Kaufman, William A. Liberti, and David L. McLean for useful discussions and feedback on the manuscript. The authors would also like to thank Dalton D. Moore, Chadbourne M. B. Smith, Gabriella Wheeler Fox, and Steven A. Redford for technical assistance.

## Author contributions

Conceptualization, E.A.D. and J.N.M.

Data curation, E.A.D.

Formal analysis, E.A.D.

Funding acquisition, E.A.D. and J.N.M

Investigation, E.A.D.

Methodology, E.A.D. and J.N.M

Resources, J.N.M.

Software, E.A.D.

Supervision, J.N.M.

Visualization, E.A.D.

Writing – original draft, E.A.D. and J.N.M

Writing – review & editing, E.A.D. and J.N.M

## Resource availability

**Lead Contact:** Requests for further information and resources should be directed to and will be fulfilled by the lead contact, Jason MacLean.

## Materials availability

This study did not generate new unique reagents.

## Data and code availability

All data reported in this paper will be shared by the lead contact upon request. All original code has been deposited on the MacLean lab Github page and is publicly available at https://github.com/MacLean-Lab-UChicago/published-code as of the date of publication. Any additional information required to reanalyze the data reported in this paper is available from the lead contact upon request.

## Supplementary Figure captions

**Supplementary Figure 1:**
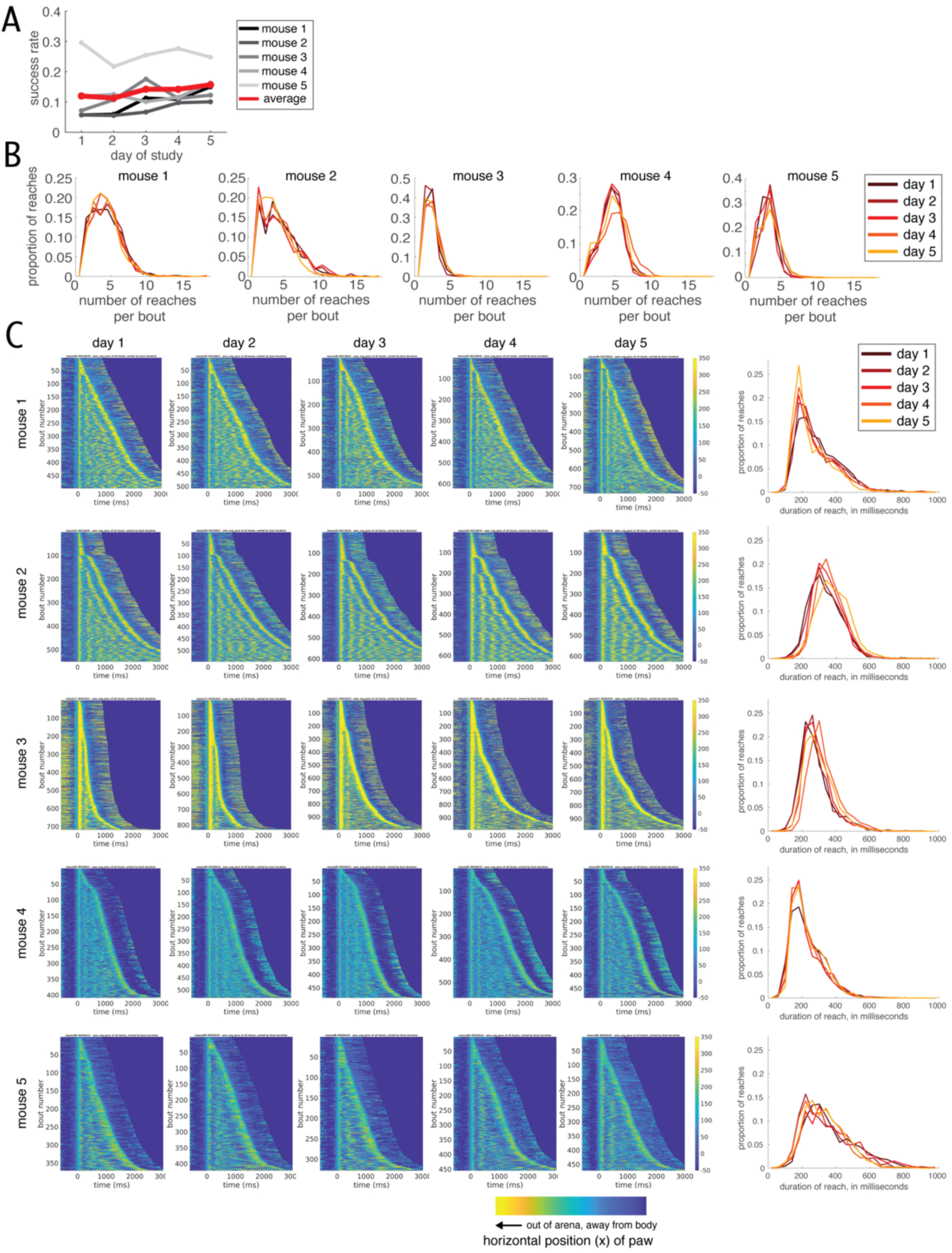
Mice exhibit stable task performance and similar bout structure over the five days. A) Success rate on each day for each animal, calculated as the proportion of reaches where the mouse successfully retrieved the seed, out of all reaches with seeds present. Average is calculated across mice. The total reach counts listed in the Results section under header *A dexterous reach-to-grasp task elicits variable yet spatiotemporally coordinated forelimb and digit movements* include reaches where a seed is not present, which are not included in success rate calculation. Thus the total number of seeds retrieved divided by the total number of reaches does not equate to the proportions shown here. See Methods for additional details about success rate calculation and criteria for including reaches in neural data analysis. B) Histograms of the number of reaches per bout, for each mouse. Color indicates day within the stability dataset. C) Same as in Fig. 2B, but for all five mice. Left, heatmaps of x position of the paw for all bouts, sorted by bout duration, using the same color scheme as Fig. 1E and 2B: yellow colors indicate further outside the arena, blue colors indicate closer to or inside the arena. Right, probability density distributions of time durations of reaches for each day. Color indicates day within the stability dataset.

**Supplementary Figure 2:**
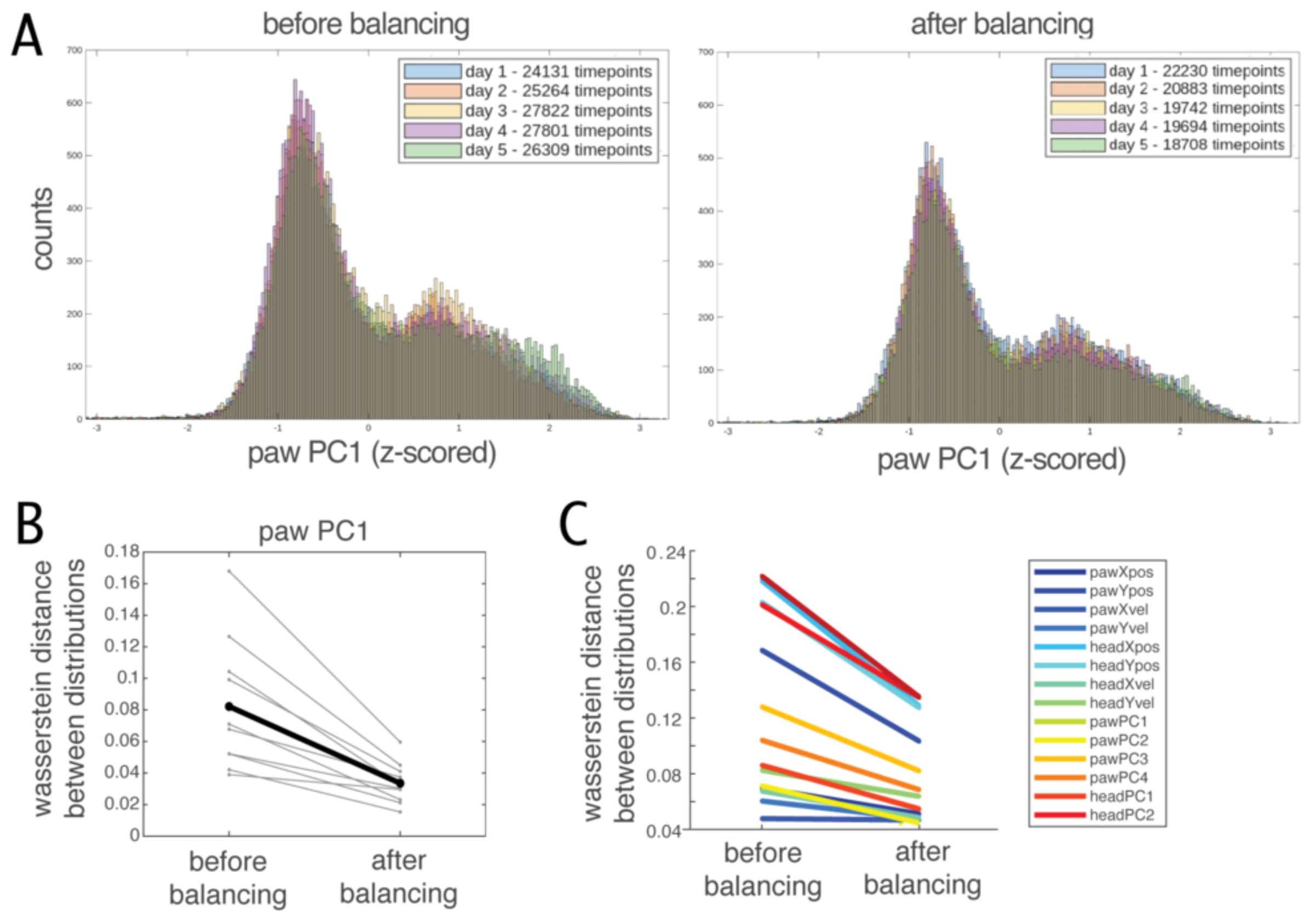
Covariate balancing increases predictor distribution similarity. A) Predictor distributions for the kinematic predictor paw PC1 (first principal components on the paw markers) for all five days for an example mouse (mouse 2). Left shows distributions before covariate balancing on the joint distribution. Right shows distributions after balancing. Color indicates day within the stability dataset. Number of timepoints in each distribution is included in the inset legend. See Methods for details of covariate balancing procedure. B) Predictor distributions for the five days become more similar to each after covariate balancing. Wasserstein distance (Earthmover’s distance) between the distributions for paw PC1 for all pairs of days for the same mouse (mouse 2), between distributions for all pairs of the five days before balancing or between distributions for all pairs of the five days after covariate balancing. Lines connect a given day-pair’s distributions’ distance before balancing with its distance after balancing. Thin gray lines indicate Wasserstein distance for individual day-pairs (5 days yields 10 day-pairs). Black line indicates mean across day-pairs. See Methods for additional details on computing Wasserstein distance. C) Grand mean of Wasserstein distances for each predictor, for the distributions before and after covariate balancing. Color indicates predictor. Grand mean is computed first across day-pairs within mouse, then across mice.

**Supplementary Figure 3:**
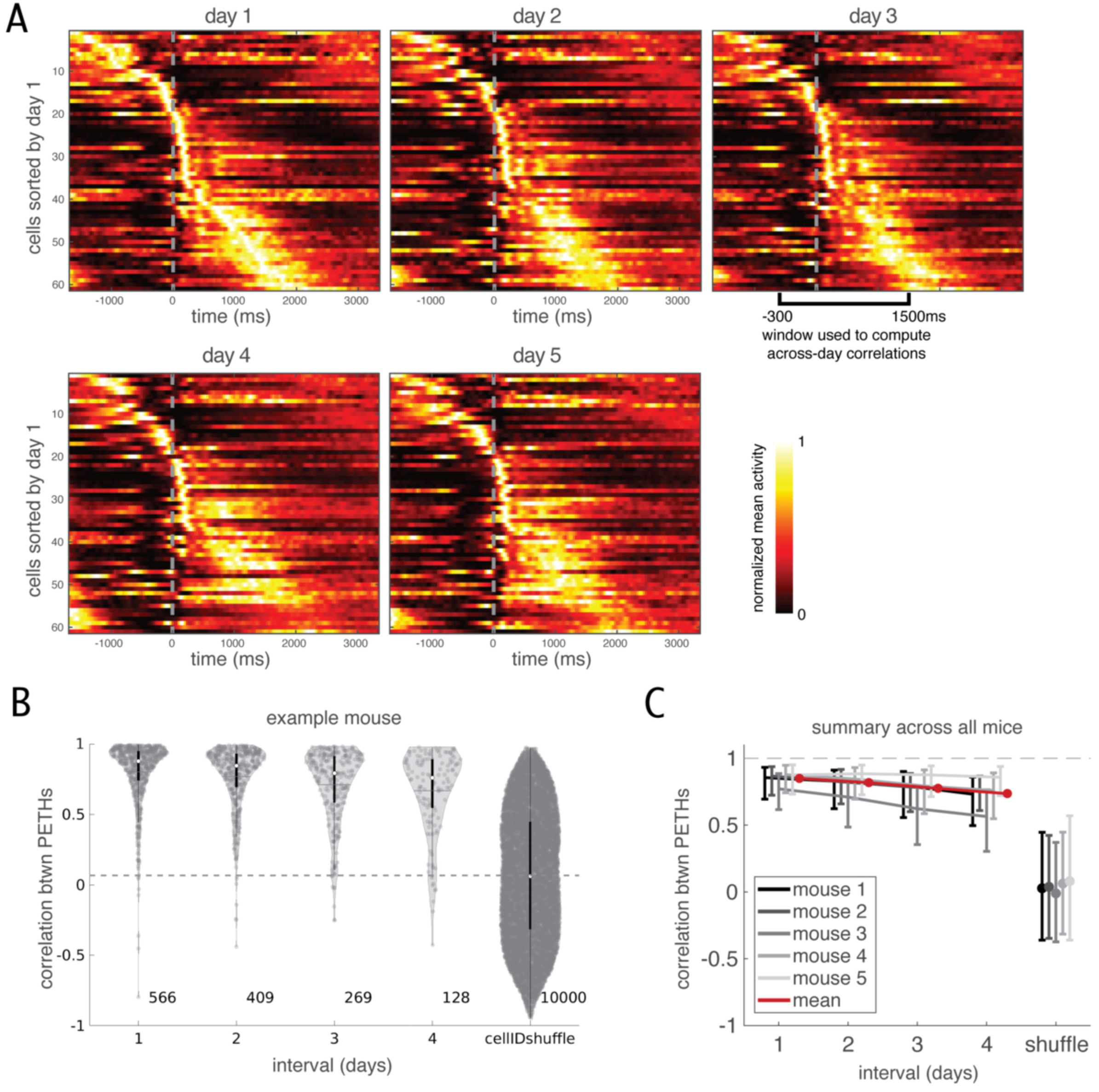
Bout-locked PETHs are also stable across days. A) Normalized peri-event time histograms (PETHs) locked to the onset of bout, for neurons found on all five days for an example mouse (mouse 4). Cells are sorted according to the time of their peak activity on either day 1. Each cell’s PETH has been normalized separately within each day within the plotted window: minimum value = 0, maximum value = 1. t = 0 indicates the time of bout onset. Onset of a bout is defined as the onset of the first reach in the bout (see Methods). B) Correlation between PETHs, as a function of the interval between the days, for all cells found on more than one day for an example mouse (mouse 4). Each datapoint is an observation of the same cell on two different days. Numbers to the bottom right of the violins indicate the number of observations included in that violin. For each violin, the white dot indicates the median of the distribution and the black box indicates interquartile range. Cell ID shuffle indicates the distribution of correlations between PETHs of two cells randomly chosen from any day (excluding any cell pairs that are registered as the same cell; see Methods). Gray dashed line indicates the median of this shuffle. C) Correlation between PETHs for all mice. Datapoint is median, error bars are interquartile range across cells for that interval for that mouse. Grayscale indicates mouse. Shuffle as in B.

**Supplementary Figure 4:**
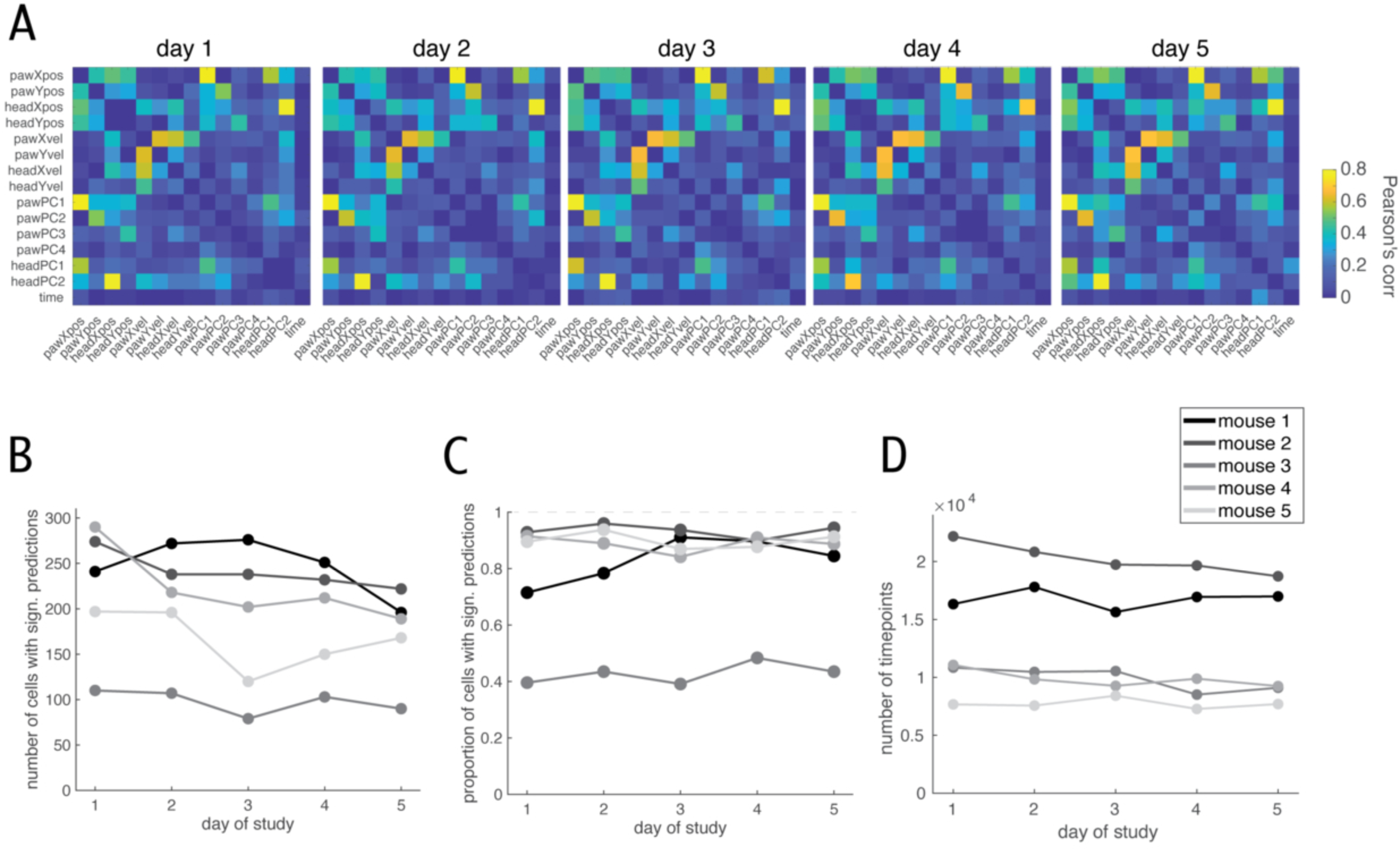
Additional details of single-trial linear models. A) Pearson’s correlation between kinematic variable predictors within each daily session, for all sessions for an example mouse (mouse 5). Correlation is computed after predictor sets have been balanced across days (see Methods for details of covariate balancing). B) Number of cells with significant linear model fits for the single-trial full model, on each day for each mouse. Grayscale indicates mouse, as in D. C) Same as in B but as proportion out of total cells imaged in the session. Grayscale indicates mouse, as in D. D) Number of timepoints from each session used to fit the linear models, after subsampling during covariate balancing (see Methods). Grayscale indicates mouse.

**Supplementary Figure 5:**
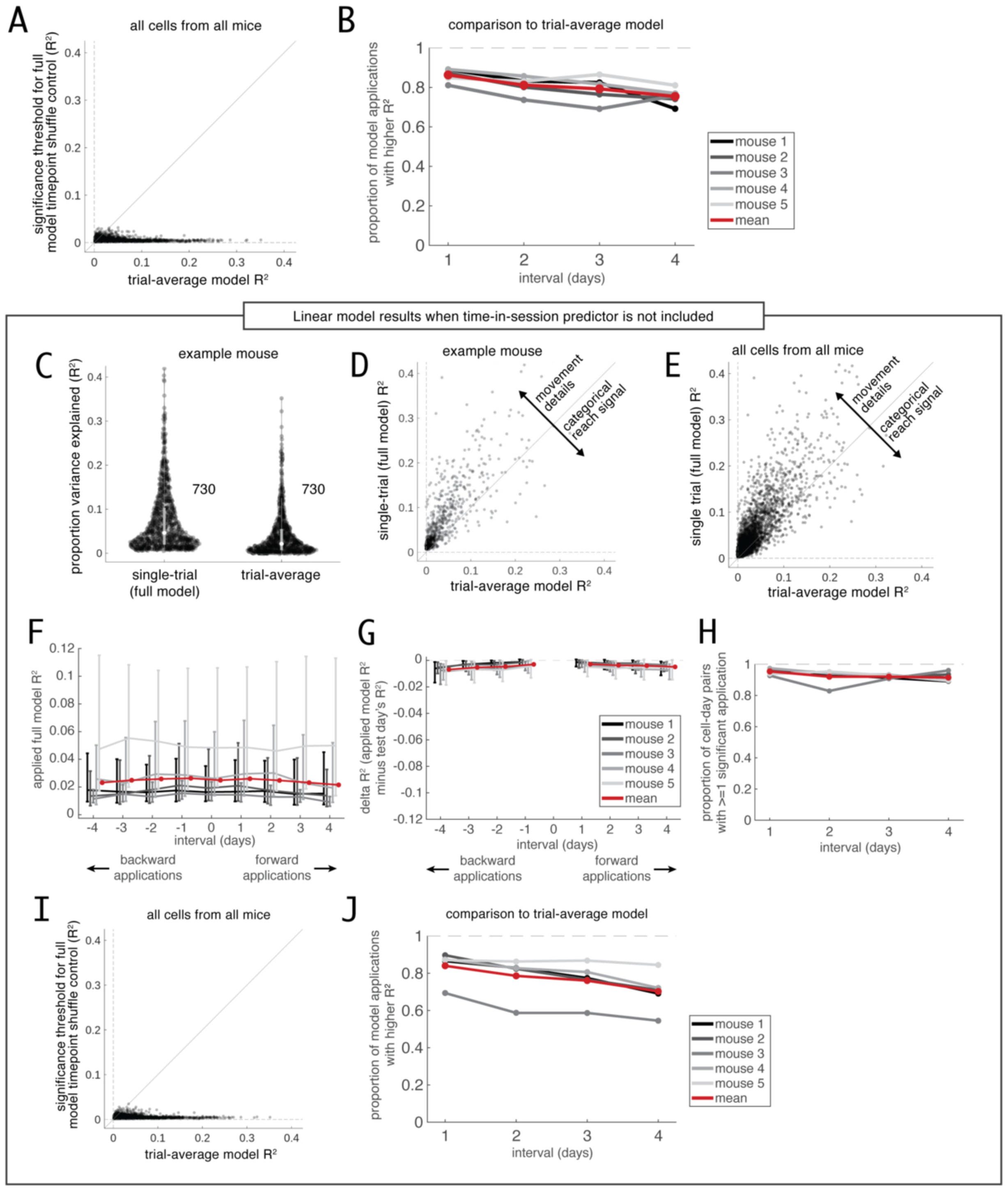
Single-trial encoding is maintained at or above the level of trial-average encoding, and all single-trial encoding results hold when models are fit without time-in-session predictor. A) For the majority of cell-days with both significant single-trial encoding and significant trial-average encoding (3295 out of 3927 cell-days, or 83.9%), the trial-average model R-squared value is higher than the significance threshold for the single-trial model (computed using a timepoint shuffle for each cell-day, see main text and Methods). Significance threshold plotted against the trial-average model R-squared value for all cell-days with a significant full single-trial model prediction and a significant trial-average model prediction, for all cells from all mice. Each datapoint corresponds to a cell on one day (cell-day), n = 3927 cell- days. B) Proportion of model applications with an applied single-trial model R-squared value higher than the test day’s trial-average model R-squared value, as a function of interval between the cell-days. Grayscale indicates mice. Only model applications for which the test day has greater single-trial encoding than trial-average encoding are included (see Results text). C) Same as Fig 5C, but for linear models fit without the time-in-session predictor. Proportion of variance explained (R-squared) by the full single-trial model or the trial-average model for all cell-days with a significant full single-trial model prediction, for an example mouse (mouse 5). Numbers to the right of each violin indicate the number of cell-days in the distribution. Each datapoint corresponds to a cell on one day (cell-day). Cells registered across days have multiple dots if they have significant full single-trial model predictions on multiple days. For each violin, the white dot indicates the median of the distribution and the gray box indicates interquartile range. D) Same as Fig 5D, but for linear models fit without the time-in-session predictor. Same data as in C, but with the full model and trial-average model R-squared values plotted against each other for each cell-day. Cells that fall above the unity line show better predictions from the single-trial model than from the trial-average model, indicating that trial-averaged movements cannot fully explain their encoding of single-trial movement details. Cells that fall below the unity line are better explained by trial-averaged movements than by single-trial movement details, indicating that their activity is best explained as a categorical reach signal that is similar across all reaches and does reflect the variability of individual reaches. E) Same as Fig 5E, but for linear models fit without the time-in-session predictor. The majority of neurons show improved predictions from the single-trial model relative to the trial-average model, indicating that they encode single-trial movement details. Same as D, but for all cell-days with significant full model predictions from all mice. F) Same as Fig 6B, but for linear models fit without the time-in-session predictor. Resulting R-squared value when a model fitted on one day is applied to another day, for all intervals between train and test days. Datapoint is median, error bars are interquartile range across train-test day pairs for that interval for that mouse. Grayscale indicates mice, as in C. Forward applications indicate the train day comes before the test day; backward applications indicate the train day comes after the test day. G) Same as Fig 6C, but for linear models fit without the time-in-session predictor. Performance of applied models relative to a test day’s own model. Difference between R-squared from an applied model and the R-squared of the test day’s own fitted model. Datapoint is median, error bars are interquartile range across train-test day pairs for that interval for that mouse. Forward and backward applications as in B. H) Same as Fig 6D, but for linear models fit without the time-in-session predictor. The majority of applied models yield statistically significant predictions. Proportion of cell-day pairs with significant model application in at least one direction, as a function of interval between the cell-days. Grayscale indicates mice, as in C. Applied models are compared to test day’s own timepoint shuffle control to determine significance (see Methods). I) Same as in A, but for linear models fit without the time-in-session predictor. J) Same as in B, but for linear models fit without the time-in-session predictor.

**Table S1.** Linear model results when time-in-session is included as a predictor, compared to when time-in- session is excluded (models are fit with only kinematic predictors)

|  | With time-in-session | Without time-in-session |
| --- | --- | --- |
| **Results related to the full single trial model fits on individual days** |  |  |
| Number of cell-days with significant full (single trial) model predictions | 4871 out of 6105 cell-days (79.8%) | 4171 out of 6105 cell-days (68.3%) |
| Proportion of cells in a session that have significant full (single trial) model predictions | 79.6% +/- 19.5%, [39.1%, 96.0%]; mean +/- st.dev. [range] | 68.0% +/- 24.0%, [19.8%, 91.1%]; mean +/- st.dev. [range] |
| Median rsq value for cell-days with significant full (single trial) predictions, median taken after pooling together all cell-days from all sessions and all mice | 0.031 | 0.021 |
| Median rsq value for all cell-days with significant full (single trial) predictions in a given session, averaged across all sessions | 0.032 +/- 0.011 [0.017, 0.060]; mean +/- st.dev. [range] | 0.024 +/- 0.012, [0.011, 0.051]; mean +/- st.dev. [range] |
| **Results related to the trial average model fits on individual days** |  |  |
| Number of cell-days with significant trial average model predictions | 4312 out of 6105 cell-days (70.6%) | 4460 out of 6105 cell-days (73.1%) |
| Proportion of cells in a session that have significant trial average model predictions | 70.4% of neurons in a given session (70.4% +/- 15.1% [31.7%, 93.6%]; mean +/- st.dev. [range]). | 73.0% +/- 14.5%, [37.8%, 93.6%]; mean +/- st.dev. [range] |
| Median rsq value for cell-days with significant trial average predictions, median taken after pooling together all cell-days from all sessions and all mice | 0.014 | 0.013 |
| Median rsq value for all cell-days with significant trial average predictions in a given session, averaged across all sessions | 0.015 +/- 0.007, [0.005, 0.030]; mean +/- st.dev. [range] | 0.014 +/- 0.007, [0.004, 0.030]; mean +/- st.dev. [range] |
| **Results related to the comparing the single trial and trial average models on individual days** |  |  |
| Number of cell-days with significant predictions for BOTH the full (single trial) model and the trial average model | 3927 | 3695 |
| Number of cell-days significantly predicted by ONLY the full (single trial) model | 944 | 476 |
| Number of cell-days significantly predicted by ONLY the trial average model | 385 | 765 |
| For cell-days significantly predicted by both the full (single-trial) model and the trial-average model, number and proportion that were better modeled by the full single-trial model than by the trial-average model | 3371 out of 3927 cell-days (85.8%);<br>85.0% +/- 7.6% of cell-days in a given session, [59.7%, 94.5%];<br>mean +/- st.dev. [range] | 2847 out of 3695 cell-days (77.1%);<br>74.1% +/- 14.1% of cell-days in a given session, [40.4%, 95.4%]<br>mean +/- st.dev. [range]) |
| **Results related to applying the full single trial models across days** |  |  |
| Number of cell-days and corresponding *pairs* of cell-days which can be considered in this analysis (aka, pairs of cell-days where both days have significant predictions for the full (single trial) model) | 4871 cell-days, corresponding to 4961 pairs of cell-days | 4171 cell-days, corresponding to 4212 pairs of cell-days |
| Number of cell-day pairs that showed significantly similar encoding in at least one direction (all between-day intervals pooled) | 4596 out of 4961 cell-day pairs (92.6%) | 3966 out of 4212 cell-day pairs (94.2%) |
| Number of cell-day pairs that showed significantly similar encoding in BOTH directions (all between-day intervals pooled) | 4283 out of 4961 cell-day pairs (86.3%) | 3684 out of 4212 cell-day pairs (87.5%) |
| Difference in delta-R-squared values for 1-day versus 4-day intervals (absolute difference between medians computed within mice, then averaged across mice, n=5 mice) | 0.004 +/- 0.002, [0.002, 0.007];<br>mean +/- st.dev. [range] | 0.0027 +/- 0.0011, [0.0013, 0.0038];<br>mean +/- st.dev. [range] |
| Number of mice for which distributions of delta-R-squared values for 1-day intervals and 4-day intervals are statistically distinguishable | 4 mice (mouse 1, 2, 3, and 4) | 3 mice (mouse 1, 2, and 4) |
| **Results related to comparing the stability of single-trial encoding to idealized stable trial-average encoding** |  |  |
| Number and proportion of cell-days (with both significant single-trial encoding and significant trial-average encoding) for which the trial average model's R-squared value exceeds the full single trial model's timepoint shuffle significance threshold | 3295 out of 3927 cell-days (83.9%) | 3215 out of 3695 cell-days (87.0%) |
| Number of cell-days and corresponding *pairs* of cell-days which can be considered for comparison to idealized stable trial-average encoding (aka, pairs of cell-days where both days have with significant predictions for BOTH the full (single trial) model and the trial average model) | 3927 cell-days, corresponding to 3833 pairs of cell-days, which is 77.3% of the 4961 pairs of cell-days originally examined with across-day linear model application. This corresponds to 7666 model applications, since each pair of cell-days has two model | 3695 cell-days, corresponding to 3638 pairs of cell-days, which is 86.4% of the 4212 pairs of cell-days originally examined with across-day linear model application. This corresponds to 7276 model applications, since each pair of cell-days |
|  | applications (one forward and one backward). | has two model applications (one forward and one backward). |
| Number and proportion of model applications for which the test day has **lower single-trial** encoding than trial-average encoding, for day-pairs where both days have significant single-trial and trial-average encoding | 1201 out of 7666 model applications (15.7%) | 1724 out of 7276 model applications (23.7%) |
| Number and proportion of model applications for which the test day has **greater single-trial** encoding than trial-average encoding, for day-pairs where both days have significant single-trial and trial-average encoding | 6465 out of 7666 model applications (84.3%) | 5552 out of 7276 model applications (76.3%) |
| For model applications for which the test day has **greater single-trial** encoding than trial-average encoding, number and proportion of model applications with an applied model R-squared value that is higher than the test day's trial-average model R-squared value (i.e. single trial encoding remains higher than idealized stable trial average encoding) | 5392 out of 6465 model applications (83.4%) | 4609 out of 5552 model applications (83.0%) |

